# Structure of Calcineurin bound to PI4KA reveals dual interface in both PI4KA and FAM126A

**DOI:** 10.1101/2024.04.09.588654

**Authors:** Alexandria L Shaw, Sushant Suresh, Matthew AH Parson, Noah J Harris, Meredith L Jenkins, Calvin K Yip, John E Burke

**Author notes:** These authors contributed equally (Alexandria L Shaw, Sushant Suresh). Lead contact: John E Burke.

## Abstract

Phosphatidylinositol 4-kinase alpha (PI4KA) maintains the PI4P and phosphatidylserine pools of the plasma membrane. A key regulator of PI4KA is its association into a complex with TTC7 and FAM126 proteins. This complex can be regulated by the CNAβ1 isoform of the phosphatase Calcineurin. We previously identified that CNAβ1 directly binds to FAM126A. Here, we report a cryo-EM structure of a truncated PI4KA complex bound to Calcineurin, revealing a direct Calcineurin interaction with PI4KA. Additional HDX-MS and computational analysis show that Calcineurin forms a complex with an evolutionarily conserved IKISVT sequence in PI4KA’s horn domain. We also characterised conserved LTLT and PSISIT Calcineurin binding sequences in the C-terminus of FAM126A. These sites are in close proximity to phosphorylation sites in the PI4KA complex, suggesting key roles of Calcineurin-regulated phosphosites in PI4KA regulation. This work reveals novel insight into how Calcineurin can regulate PI4KA activity.

## Introduction

Phosphoinositide lipids act as master second messengers which regulate myriad cellular processes ^1–3^. One of the most abundant phosphoinositides is phosphatidylinositol 4-phosphate (PI4P), with its concentration varying by cell-type, at ∼1% of total cellular phospholipid ^4^. PI4P is found in the cell in various subcellular membrane locations, including the Golgi, the endo-lysosomal system, and the plasma membrane (PM) ^4^. PI4P is generated by the action of four different phosphatidylinositol 4-kinases ^5,6^, with the plasma membrane pool being primarily generated by type III phosphatidylinositol 4-kinase alpha (PI4KIIIα, which will be referred to by its gene name *PI4KA* for the remainder of the manuscript). Given the low abundance of PI at the PM ^7,8^, the generation of PI4P by PI4KA at the plasma membrane is essential for the generation of membrane asymmetry, partially through its critical role in controlling phosphatidylserine (PS) transport to the plasma membrane through ORP5/8 ^9^. This function of PI4KA is evolutionarily conserved from yeast to humans.

PI4KA is a multi-domain protein, composed of an alpha-solenoid (horn) domain, a dimerization domain, a helical domain, and a bi-lobal kinase domain ^10^. It is functional as a large multi-protein complex, primarily associated with two scaffolding proteins, TTC7 (with two possible isoforms TTC7A or TTC7B) and FAM126 (with two possible isoforms FAM126A or FAM126B) ^10–12^. PI4KA and TTC7 are highly conserved throughout all eukaryotes, with FAM126 being absent in many simple unicellular eukaryotes ^13,14^. The PI4KA-TTC7-FAM126 complex forms a dimer of trimers, with TTC7 and FAM126 stabilizing PI4KA, which together will be referred to as the PI4KA complex for the remainder of the manuscript ^10–12^. Functionally, PI4KA is recruited to the membrane by a palmitoylated EFR3 protein (with two possible isoforms EFR3A/EFR3B ^15^. The exact binding site of EFR3 to the PI4KA complex is not fully resolved but putatively involves an evolutionarily conserved C-terminal region of TTC7 ^16^. PI4KA faces a complicated challenge in the generation of PI4P at the plasma membrane, as its substrate PI is present at only trace levels ^7,8^, with the exact mechanism of how PI4KA is regulated and activated at the plasma membrane still not fully understood.

PI4KA, TTC7, FAM126, and its accessory proteins are implicated in myriad human diseases. Intriguingly, both the loss-of-function and gain-of-function of PI4KA can be drivers of human disease ^17^. Down-regulation of PI4KA signaling is observed in intestinal atresia driven by mutations in TTC7A ^18^, hypomyelination and congenital cataracts by a deficiency in FAM126A ^19^, and neurological, intestinal and immunological diseases caused by loss-of-function mutations in PI4KA ^20,21^.

One of the recently discovered regulators of PI4KA signaling is the protein phosphatase Calcineurin, which is a ubiquitously expressed serine/threonine phosphatase activated by Ca^2+^ signaling ^22^. Calcineurin is composed of two subunits, calcineurin A (CNA) and calcineurin B (CNB), with multiple isoforms of each subunit. We have found that the lipidated CNAβ1 isoform of Calcineurin is a direct regulator of PI4KA signaling, and upon palmitoylation is recruited to the PM ^23^. Cellular studies have shown that Calcineurin inhibition causes reduced PI4P synthesis following hormone-mediated PI(4,5)P2 depletion, and we identified a short linear PxIxIT motif (SLiM) in FAM126A that mediates binding to CNAβ1 ^23^.

To fully explore the molecular mechanisms of how Calcineurin can bind to and regulate the PI4KA complex we have utilised a synergistic application of cryo-EM, hydrogen deuterium exchange mass spectrometry (HDX-MS), and AlphaFold3 modelling. Our cryo-EM structure reveals that Calcineurin not only binds the FAM126A disordered tail but also forms an evolutionarily conserved interface with the horn region of PI4KA. We used AlphaFold3 modelling to predict how PI4KA interacts with Calcineurin, with the predicted interface validated experimentally by HDX-MS. This map also had increased local resolution in the horn compared to previous cryo-EM analysis allowing for the building of a complete model of the PI4KA horn, as well as the interface between TTC7B and the N-terminus of the PI4KA horn. HDX-MS also revealed a Calcineurin binding site in a disordered loop in PI4KA’s dimerization domain, which could interact with CNB’s LxVP binding motif. These evolutionarily conserved PI4KA Calcineurin SLiM motifs orient the active site of Calcineurin near multiple evolutionarily conserved phosphorylation sites of unknown function. Overall, our findings provide useful insight into the molecular and structural basis of PI4KA complex regulation by Calcineurin.

## Results and Discussion

### Cryo-EM structure of the PI4KA-Calcineurin complex

We purified the human PI4KA complex with a truncated FAM126A tail (full-length PI4KA/TTC7B and FAM126A ΔC (aa 1-308), referred to in the rest of the manuscript as PI4KA complex (Fig 1A). This construct lacks the FAM126A C-terminus, which was previously identified as a binding site for Calcineurin (composed of a dimer of CNA and CNB subunits). We had previously observed a change in protein dynamics in the horn region of PI4KA in full-length PI4KA-TTC7B-FAM126A upon binding Calcineurin ^23^, and it was unknown if this was driven by allosteric conformational changes or acted as a direct Calcineurin binding interface. To further investigate this, we purified the Calcineurin heterodimer (alpha isoform of CNA with CNB) to conduct structural studies of the PI4KA- Calcineurin interaction. This was done as all Calcineurin isoforms interact with their substrates through the same mechanism(s) and the alpha isoform of CNA is best behaved in solution and therefore more suitable for *in vitro* structural studies.

**Figure 1.**
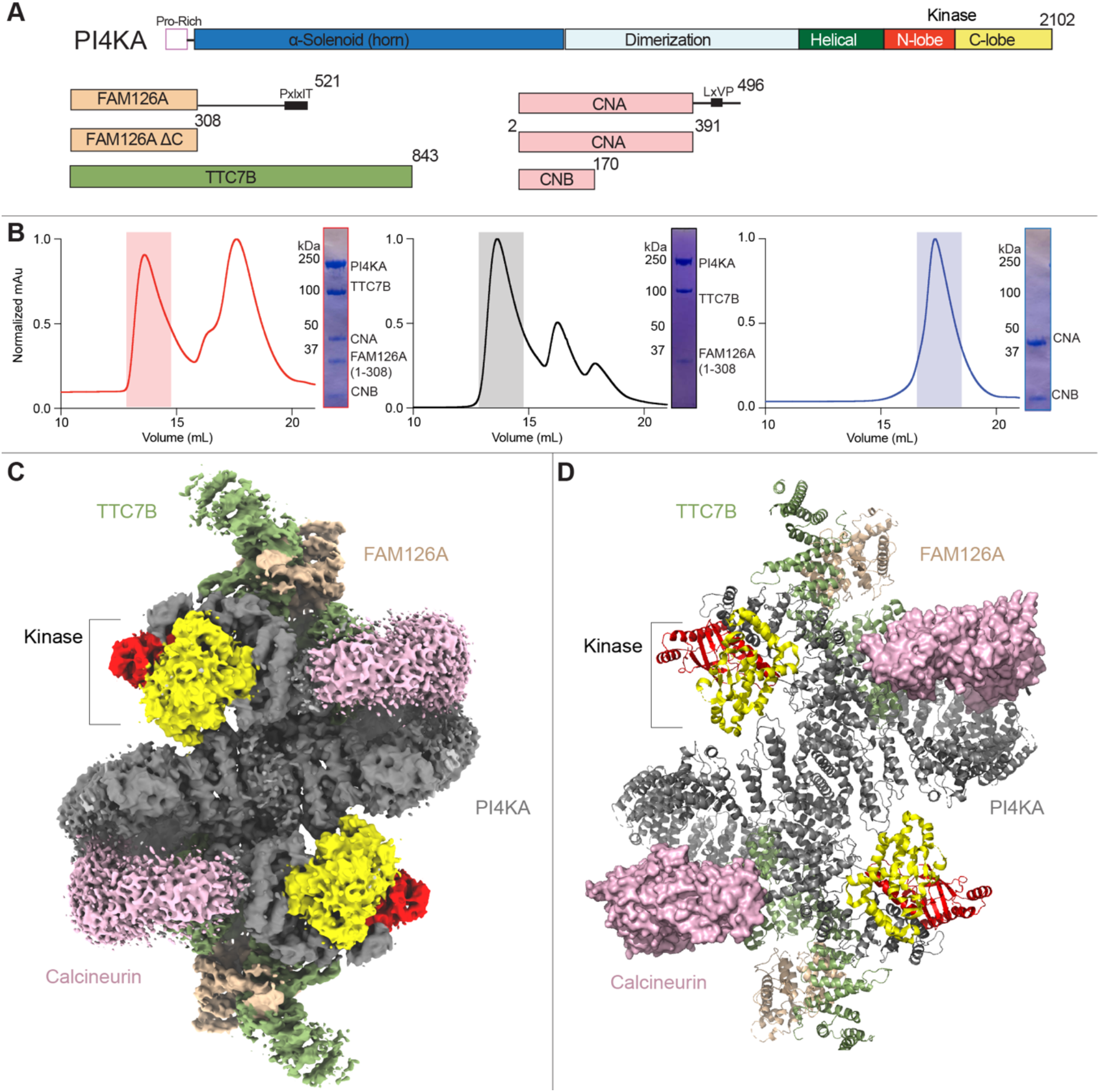
Calcineurin forms a complex with the PI4KA complex (see supplemental figures 1-3) **A.** Domain schematics of catalytic PI4KA, regulatory proteins TTC7B and FAM126A (Full length and truncated FAM126A ΔC used in all experiments) and the serine/threonine phosphatase Calcineurin (heterodimer of CNA (top: Full length β1 isoform, bottom: truncated ⍺1 isoform used in all experiments), and CNB). **B.** Gel filtration elution profiles of PI4KA complex apo (black), Calcineurin apo (blue), or PI4KA complex bound to Calcineurin (red). SDS-polyacrylamide gel electrophoresis image of specified peaks. **C.** Cryo-EM density map of the PI4KA complex bound to Calcineurin. **D.** Cartoon/surface representation of the PI4KA complex bound to Calcineurin modeled from the Cryo-EM map coloured according to the in-figure text legend.

To validate if the PI4KA complex could directly interact with Calcineurin in the absence of the C-terminus of FAM126A, we analysed three complexes by gel filtration: PI4KA complex, Calcineurin, and the co-complex. Elution profiles were consistent with the formation of a stable stochiometric dimer of pentamers (pentamer composed of PI4KA, TTC7B, FAM126A ΔC, CNA, and CNB, Fig. 1B). To delineate the molecular basis for how Calcineurin binds to PI4KA we examined the architecture using an approach combining cryo-EM, HDX-MS, and AlphaFold3 modelling.

We first conducted cryo-EM analysis of the PI4KA complex bound to Calcineurin. Although negative stain analysis revealed that the purified PI4KA complex bound to Calcineurin was homogeneous, our initial attempts to obtain good 3D reconstructions were unsuccessful due to preferred particle orientation. We reasoned that this could likely be due to the dynamic nature of Calcineurin relative to PI4KA, and PI4KA’s proposed membrane binding interface leading to the accumulation of PI4KA at the air-water interface. To obtain a more “rigid” complex for cryo-EM analysis, we treated the PI4KA complex bound to Calcineurin with BS^3^ crosslinker, followed by gel filtration to remove aggregate proteins. The crosslinked complex eluted from gel filtration equivalently to the non-crosslinked complex. Analysis of this complex allowed for the refinement of 235,760 particles leading to a 3D reconstruction at a nominal resolution of 3.50 Å (Fig 1C, Fig S1). The density map was of sufficient quality to allow for automated and manual building of the majority of the PI4KA subunit, with a clear orientation of the available cryo-EM and X-ray crystallography models of TTC7B-FAM126A in the electron density, although we had only weak density for the N-terminal region of TTC7B therefore we removed this from our molecular model. ^10,16^. The regions with the lowest local resolution correspond to the two Calcineurin molecules. Despite this, the low local resolution of the map for Calcineurin (∼7 Å) was sufficient to allow for a rigid body fit of previously solved high resolution Calcineurin structures into the density, and allowed for an orientation of Calcineurin relative to PI4KA. However, at this resolution we could not experimentally define the residues mediating the interaction between PI4KA and Calcineurin, and this interface was predicted using AlphaFold3, and validated using HDX-MS (described below).

Our cryo-EM map captures more details in PI4KA than compared to the previously reported cryo-EM structure of the PI4KA complex ^10^. While the interface between TTC7 and PI4KA in the dimerization and helical domains of PI4KA have been extensively characterised ^10^, the interaction between the solenoid domain of PI4KA and Calcineurin have been structurally unresolved. Using a combination of manual model building and AlphaFold/trRosetta modelling of specific regions ^24,25^ we were able to confidently build an additional ∼700 residues of PI4KA corresponding to the solenoid α helices in the horn domain (Fig S2 and S3). Intriguingly, the solenoid horn domain showed a slight deviation compared to the previous cryo-EM structure of PI4KA (Fig S2B). It is not known if this conformational difference was driven by the interaction with Calcineurin or by the chemical crosslinker used to stabilize the assembly. The local resolution of the contact between the N-terminus of PI4KA and TTC7B allowed for unambiguous determination of the residues mediating this interface (Fig 2, S1C, S2C). The interface between the N-terminus of the horn domain (composed of solenoid α helices 1+2 [residues 31-59], Fig 2C) and the C-terminus of TTC7B was evolutionarily conserved through *C. elegans* (Fig 2D), while only weakly conserved in yeast. It has been previously shown that disease-linked C-terminal truncations in TTC7A that disrupt this interface can still bind to PI4KA, although they show decreased PI4KA activity ^10^. Studies on PI4KA were initially complicated by improper annotation of the N-terminal start site (ORF starting at M59) ^15^, with our results revealing the molecular basis for why the two evolutionarily conserved N-terminal PI4KA helices immediately before this site are critical in mediating contact with the C-terminus of TTC7. This further reinforces that any cellular studies examining PI4KA’s roles with a construct containing the improperly annotated start site should be interpreted with extreme caution.

**Figure 2.**
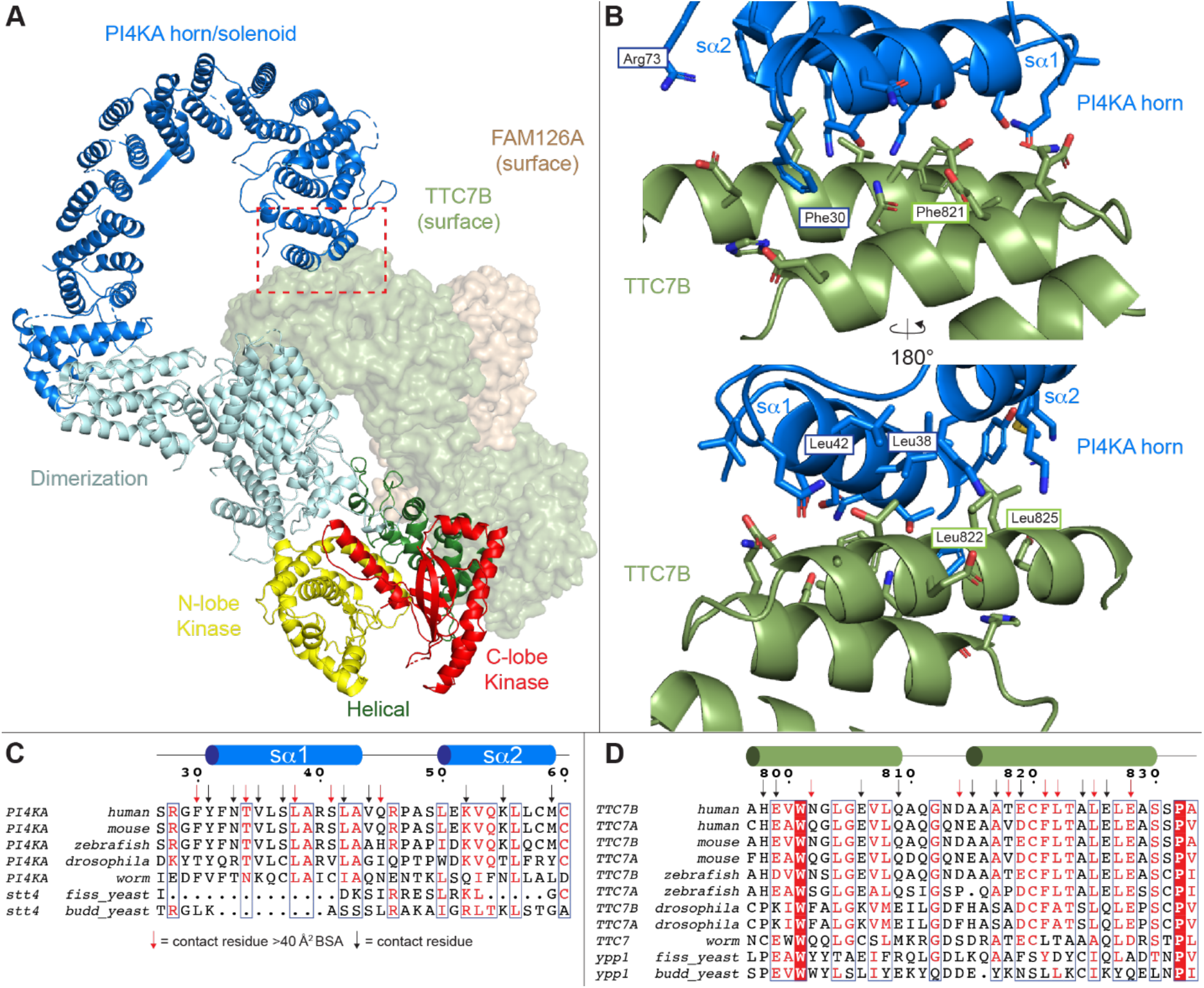
The N-terminus of the PI4KA solenoid interacts with the TTC7B C-terminal region. A. Cartoon representation of an isolated PI4KA/TTC7B/FAM126A trimer. PI4KA is shown as a cartoon with individual domains coloured according to the in-figure text. TTC7B and FAM126A are shown as surfaces coloured according to the in-figure text. Red box depicts the PI4KA solenoid-TTC7B interface. **B.** Cartoon representation of the PI4KA solenoid-TTC7B interaction interface. Side chains are shown for residues within 6 Å of either interface, which are either annotated on the cartoon or in panel C/D. Annotated residues on the cartoon were chosen to provide readers with a point of reference. **C.** Multiple sequence alignment of PI4KA from *Homo sapiens, Mus musculus, Danio rerio, Drosophila melanogaster, Caenorhabditis elegans*, *Schizosaccharomyces pombe,* and *Saccharomyces cerevisiae.* s⍺1 and s⍺2 are annotated above the alignment. Residues within 6 Å shown in panel B are annotated using arrows, with red arrows indicating a BSA of > 40 Å^2^. **D.** Multiple sequence alignment of TTC7A/B from *Homo sapiens, Mus musculus, Danio rerio, Drosophila melanogaster, Caenorhabditis elegans*, *Schizosaccharomyces pombe,* and *Saccharomyces cerevisiae.* TTC7B secondary structure observed from experimental data is annotated above the alignment. Residues within 6 Å shown in panel B are annotated using arrows, with red arrows indicating a BSA of > 40 Å^2^.

### Cryo-EM, AlphaFold3, and HDX-MS analysis of PI4KA-Calcineurin binding site

The local resolution of Calcineurin allowed us to fit previously solved structures of Calcineurin into the density ^26^, however, it was not sufficient to model the exact PI4KA residues mediating binding. To precisely map the location of the Calcineurin binding site we utilised AlphaFold3 to predict the interface, and HDX-MS, which is a technique for probing protein dynamics and characterizing protein-protein interfaces ^27–29^ to validate the prediction.

Having experimentally identified a binding event between Calcineurin and PI4KA, we carried out AlphaFold3 analysis to predict the interface responsible for this interaction^30^. We carried out searches composed of Calcineurin (CNA alpha 1 (2-391) and CNB) along with our PI4KA complex construct (PI4KA/TTC7B/FAM126A 1-308). The resulting prediction had overall ptm and iptm scores of 0.76 and 0.74, respectively, consistent with a confident multi-protein prediction (see methods). AlphaFold3 predicted with high confidence a beta strand (residues 536-541 containing the IKISVT sequence) in PI4KA binding to the NIR region of Calcineurin that mediates binding to PxIxIT motifs, with predicted alignment error (PAE) scores consistent with a stable interface (Fig. 3A/B/C). We also observed that an unstructured loop in PI4KA’s dimerization domain formed an interaction with CNB at the LxVP SLiM motif binding pocket, however, with lower pLDDT and PAE scores compared to the PI4KA-CNA interface. There were no other predicted interactions between PI4KA and Calcineurin. The AlphaFold3 prediction fit well into our low-resolution cryo-EM map after rigid body refinement.

**Figure 3.**
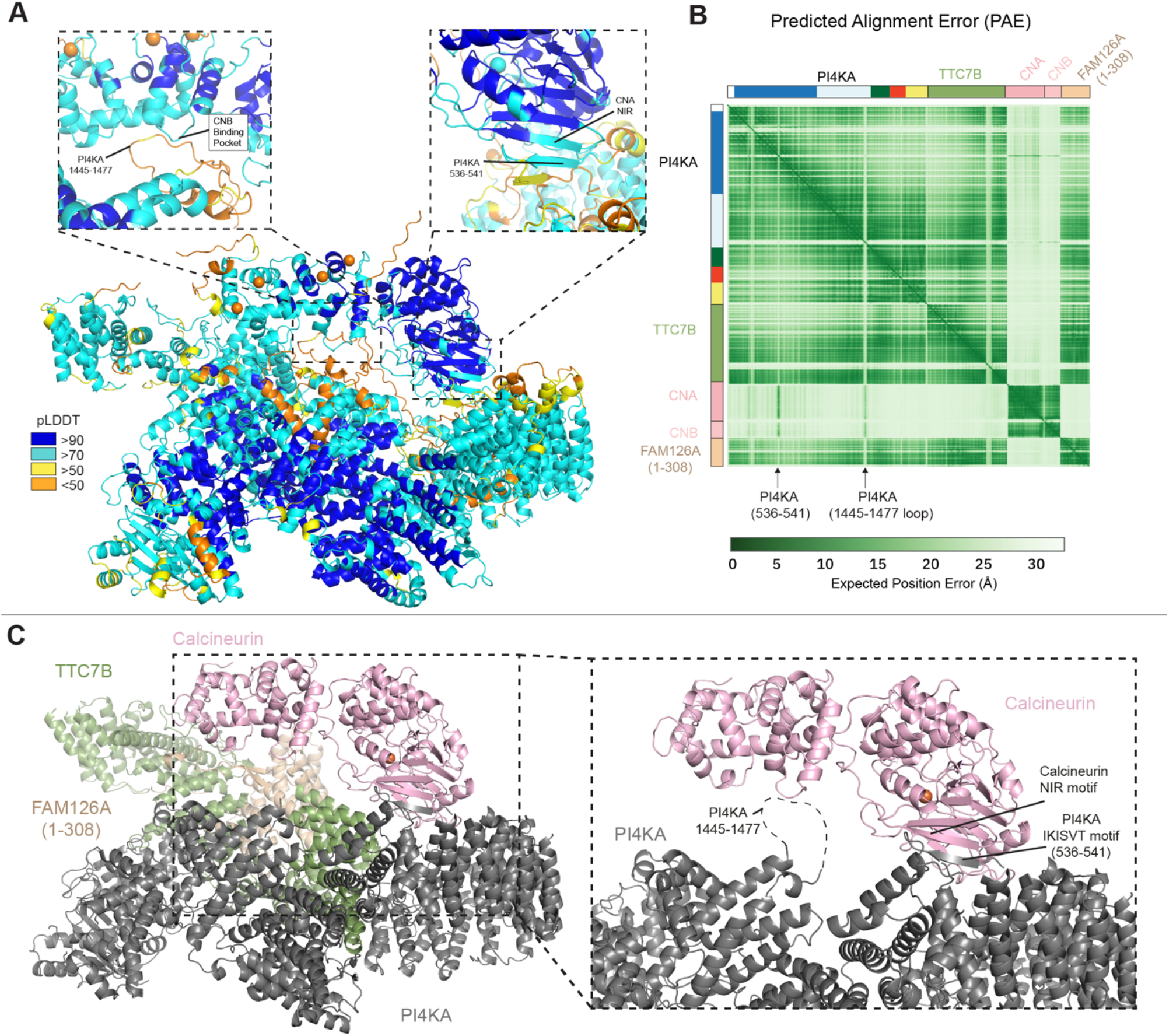
AlphaFold3 predicts a dual interface between PI4KA and Calcineurin. A. AlphaFold3 prediction of PI4KA/TTC7B/FAM126A ΔC in complex with Calcineurin and co-factors with the per-residue confidence metric predicted local-distance difference test shown as per the legend, with zoom-ins of the predicted PI4KA interface with (L) the Calcineurin B LxVP binding pocket and (R) Calcineurin A NIR motif. **B.** Predicted aligned error (PAE) of the AlphaFold3 prediction of the PI4KA/TTC7B/FAM126A (1-308) in complex with Calcineurin and co-factors. **C.** AlphaFold3 prediction of PI4KA/TTC7B/FAM126A ΔC in complex with Calcineurin and co-factors, with pLDDT <60 removed. (R) Zoom in on the predicted PI4KA – Calcineurin interface, with TTC7B and FAM126A ΔC removed. Dotted grey line represents region with a low pLDDT score.

To validate the AlphaFold3 prediction, HDX-MS experiments were conducted. The rate at which deuterium is exchanged on the amide backbone is dependent on the stability of protein secondary structure. Deuterium incorporation is localised at peptide-level resolution through the generation of pepsin-generated peptides, therefore allowing us to map the protein-protein interface at peptide level resolution. HDX experiments compared the dynamics of Calcineurin and the PI4KA complex both free in solution and when in the co-complex. Initial experiments were carried out at a high concentration of PI4KA complex and Calcineurin (1.3 µM final) at a 1:1 ratio complex, with deuterium incorporation measured over three time points (3, 30, 300 s at 18°C at pH 7.0). These experiments were carried out under three conditions: Calcineurin apo, PI4KA complex apo, and the co-complex, allowing us to map changes on both sides of the interface. These experiments revealed that the interacting motifs in PI4KA were all in regions that were disordered ^23^, consistent with the biophysical properties of the motifs that mediate interaction(s) with Calcineurin ^31,32^. To better cover the deuterium incorporation of these highly dynamic regions we carried out an additional experiment using a pH and temperature regime optimized for disordered regions (pH 6.5, 0°C) ^33^. Experiments were carried out with a 1:2 ratio of the PI4KA complex to Calcineurin (1.25 and 2.5 µM final), with deuterium incorporation measured over three time points (3, 10, 30 s). The full raw deuterium incorporation data for all HDX-MS experiments are provided in the source data. Coverage of all proteins was over 85%, with all HDX-MS processing statistics available in the source data.

We observed a significant decrease (significant change defined as greater than both 5% and 0.45 Da at any time point in any peptide with an unpaired two-tailed t test p<0.01) in deuterium exchange at residues 331-347 (NIR binding site) in CNA, residues 119-129 (LxVP binding pocket) in CNB, and residues 538-549 and 1417-1438 in PI4KA. The 538-549 region is an unstructured region between solenoid α helices 17 and 18 (Fig 4A-C, S4). This is similar to decreases in exchange observed when using full length PI4KA complex ^23^, however the absence of the FAM126A C-terminus allows for a confident validation of a protein-protein interface. The additional decrease in exchange was observed in PI4KA at residues 1417-1438, which covers an unstructured region containing sequence previously observed to bind CNB ^34^.

**Figure 4.**
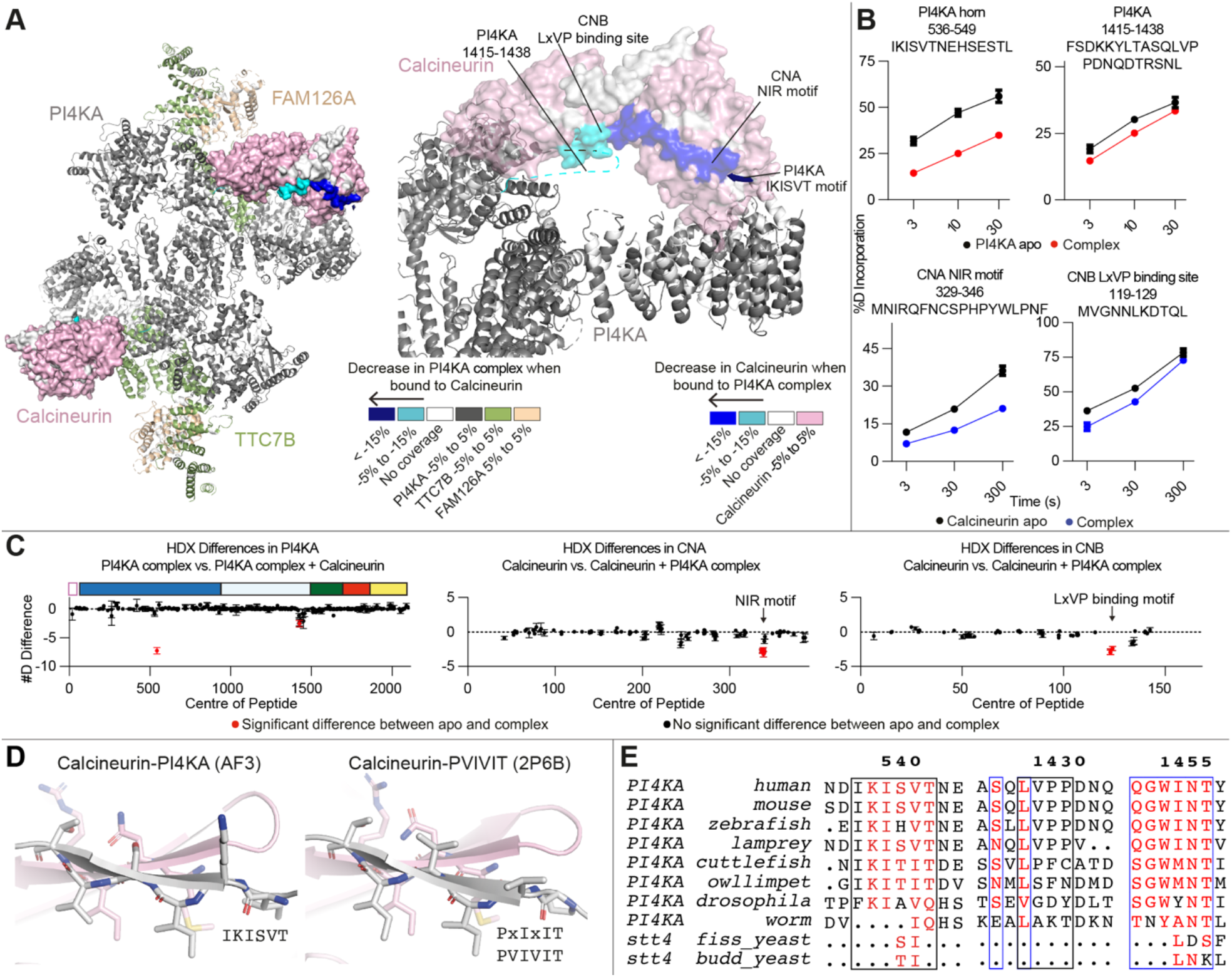
Calcineurin interacts with the PI4KA solenoid. A. (L) PI4KA complex (pH 6.5, 0°C) and Calcineurin (pH 7.0, 18°C) peptides showing significant differences in deuterium exchange (defined as >5%, 0.45 Da, and p<0.01 in an unpaired two-tailed t test at any time point) upon complex formation. Differences are mapped on the structural model, with a disordered loop showing changes in HDX being represented by a dotted line. Differences are indicated by the legend. (R) Differences are mapped on a zoomed in structural model of the interface. Differences are indicated by the legend in A. **B.** Selected deuterium exchange time courses of PI4KA (pH 6.5, 0°C) and Calcineurin (pH 7.0, 18°C) peptides that showed significant decreases in exchange upon complex formation. Error is shown as standard deviation (SD) (n=3). **C.** Sum of the number of deuteron difference of (L) PI4KA upon complex formation with Calcineurin (pH 6.5, 0°C), (M) Calcineurin A or (R) Calcineurin B upon complex formation with the PI4KA complex (pH 7.0,18°C), analysed over the entire deuterium exchange time course for PI4KA complex/Calcineurin. (M/R) Each point is representative of the centre residue of an individual peptide. Peptides that met the significance criteria described in panel D are coloured red. Error is shown as the sum of standard deviations across all 3 time points (SD) (n=3). **D.** Cartoon representation of the (L) IKISVT mediated interaction with Calcineurin’s NIR motif and (R) canonical PxIxIT mediated interaction with Calcineurin’s NIR motif. **E.** Multiple sequence alignment of PI4KA from *Homo sapiens, Mus musculus, Danio rerio, Petromyzon marinus, Sepia pharaonis, Lottia gigantea, Drosophila melanogaster, Caenorhabditis elegans*, *Schizosaccharomyces pombe,* and *Saccharomyces cerevisiae,* showing conservation of the PI4KA IKISVT motif, and regions of the PI4KA dimerization domain unstructured loop.

We conducted multiple sequence alignments of PI4KA to identify conserved elements that may mediate binding. We found that the IKISVT binding site for Calcineurin in PI4KA and the potential LVPP binding site showing decreased exchange by HDX-MS were evolutionarily conserved throughout all vertebrates, with them being partially conserved in chordates, and lost in arthropods (Fig. 4E). However, we found that an additional WINT motif in the unstructured 1427-1477 loop was also evolutionarily conserved throughout all chordates and lost in arthropods (Figure 4E), with this binding site being predicted by AlphaFold3. The binding of IKISVT bound to the NIR region of Calcineurin is consistent with Calcineurin binding to the canonical PxIxIT docking groove (Fig. 4D), and is also consistent with the high degree of plasticity observed in Calcineurin SLiM motifs ^31,32^. To fully understand the role of either the LVPP or WINT sequences in the 1427-1477 PI4KA loop binding to CNB’s LxVP binding pocket, further experiments will be required.

### Comparison of the FAM126A-Calcineurin and PI4KA-Calcineurin binding sites

To fully understand the myriad molecular interactions mediating how the PI4KA complex interacts with Calcineurin, we wanted to determine how the C-terminus of FAM126A, which is absent from our cryo-EM structure, might interact with Calcineurin. To investigate this interaction, we purified the FAM126A C-terminus [FAM126A (415- 521)] and used HDX-MS to compare the dynamics of FAM126A (415-521) apo to FAM126A (415-521) in complex with Calcineurin (Fig. 5, S4). The full raw deuterium incorporation data for all HDX-MS experiments are provided in the Source Data.

**Figure 5.**
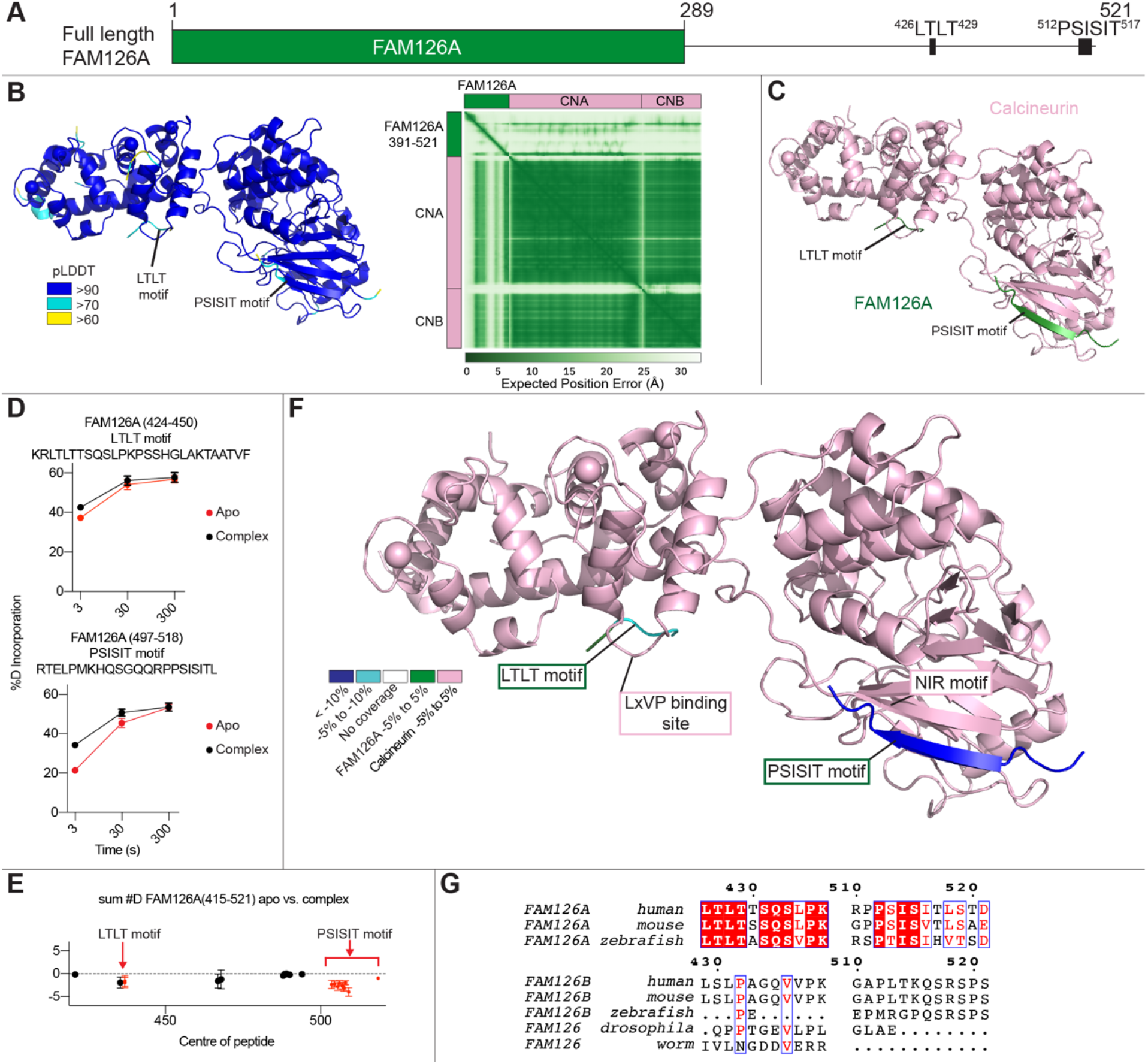
FAM126A LTLT motif interacts with CNB’s LxVP binding site. A. Domain schematic of FAM126A highlighting the disordered C-terminal which contains conserved LTLT and PSISIT motifs. **B.** (L) AlphaFold3 prediction of Calcineurin in complex with the FAM126A C-terminal tail coloured with the per-residue confidence metric predicted local-distance difference test (pLDDT) as per the legend, and pLDDT <60 removed. (R) Predicted aligned error (PAE) of the AlphaFold3 search of Calcineurin and the FAM126A C-terminal tail. **C.** AlphaFold3 prediction of Calcineurin in complex with the FAM126A C-terminal tail with pLDDT <60 removed. Domains are coloured according to the in-figure text. **D.** Selected deuterium exchange time courses of FAM126A (415-521) and Calcineurin peptides that showed significant (defined as >5% 0.45 Da, and p<0.01 in an unpaired two-tailed t test at any time point) decreases in exchange. Error is shown as standard deviation (SD) (n=3). **E.** Sum of the number of deuteron difference of FAM126A (415-521) upon complex formation with Calcineurin analysed over the entire deuterium exchange time course for FAM126A (415-521). Each point is representative of the centre residue of an individual peptide. Peptides that met the significance criteria described in F are coloured red. Error is shown as the sum of standard deviations across all 3 time points (SD) (n=3). **F.** Peptides showing significant differences in deuterium exchange upon complex formation. Differences are mapped on the AlphaFold3 prediction shown in panel C. Differences are indicated by the legend. **G.** Multiple sequence alignment of FAM126A (from *Homo sapiens, Mus musculus,* and *Danio rerio*) and FAM126B/FAM126 (from *Homo sapiens, Mus musculus, Danio rerio, Drosophila melanogaster,* and *Caenorhabditis elegans*.

The HDX-MS experiments were carried out at a 1:1 ratio, with deuterium incorporation measured over three time points (3, 30, 300 s at 0℃). As the C-terminus of FAM126A is intrinsically disordered in the absence of binding partners (Fig 5A), we carried out HDX-MS experiments at a temperature of 0℃ and a pH of 6.5 to optimize the experiment for rapidly exchanging amide hydrogens ^33^. Two areas in FAM126A (415-521) had significant changes in deuterium exchange upon complex formation with Calcineurin (significant change defined as greater than both 5% and 0.45 Da at any time point in any peptide with an unpaired two-tailed t test p<0.01). As expected, the largest significant decrease in deuterium exchange occurred in peptides containing the PSISIT binding site (494-521) (Fig. 5D-F). Another site with a significant decrease in deuterium contained an LTLT sequence (423-450), which we postulated may be able to bind in the LxVP binding pocket ^35^. Alphafold3 was used to predict the structure of the FAM126A-mediated interaction of PI4KA and Calcineurin (Fig. 5B-C) ^30^, with the LTLT (426-429) and PSISIT (512-517) sequences predicted to bind in the LxVP and PxIxIT binding sites on Calcineurin, with predicted alignment error indicative of a stable interface.

## Discussion

We had previously discovered that PI4KA was regulated by the plasma membrane localised CNAβ1 isoform of Calcineurin, with Calcineurin playing a role in regulating PI4P synthesis at the plasma membrane downstream of GPCR signaling ^23^. In this study, we aimed to investigate the full complement of molecular mechanisms underlying this regulatory role of Calcineurin in PI4KA signaling. Our synergistic application of cryo-EM, HDX-MS, and AlphaFold3 modelling has established that the PI4KA complex contains two proteins that have evolutionarily conserved and distinct Calcineurin binding sites. PI4KA has two regions that bind Calcineurin, one located in the horn domain, and one in the dimerization domain, with FAM126A binding to Calcineurin through its disordered C-terminus.

An important question posed from this work, is the evolutionary mechanism explaining why PI4KA has some of the largest variations in Calcineurin binding SLiM motifs ^31,32^. This could be driven by CNAβ1 being the only Calcineurin isoform to be directly localised to the plasma membrane through palmitoylation with it being at a higher local concentration relative to PI4KA compared to other calcineurin substrates through membrane co-localization ^23^. However, variation from the canonical LxVP and PxIxIT motifs in Calcineurin regulated proteins has been observed, including Calcineurin binding proteins containing IAIIIT ^36^ and LVPP sequences at their binding sites ^34^. Another important unanswered question of our work is why the PI4KA complex has evolved to have Calcineurin binding sites on both FAM126A and PI4KA. This is especially puzzling as PI4KA can bind either FAM126A or FAM126B, with only FAM126A containing a SLiM motif. Further cellular and organismal analysis of mutants in both the PI4KA and FAM126A sites will be required to understand their precise role in PI4KA signaling, and any FAM126A-specific functions in Calcineurin/PI4KA signaling.

Canonically, Calcineurin binds to substrate SLiM motifs to orient substrate phosphorylation sites towards the phosphatase active site of Calcineurin ^22,31,32^. We previously identified a CNAβ1 regulated phosphosite in the C-terminus of FAM126A ^23^, which suggested that there likely could be additional phosphorylation sites in the vicinity of the PI4KA horn site. There is an annotated phosphorylation site at S1436 near a region with decreased deuterium exchange in PI4KA, which could be dephosphorylated by Calcineurin. There are also extensive phosphorylation sites present in the C-terminus of both TTC7A and TTC7B that are in close spatial proximity to Calcineurin (Fig. S5A-C) ^37^. These sites are present in a disordered loop (642-703 in TTC7B, and 650-688 in TTC7A) that could be accessible to the active site of Calcineurin when comparing AlphaFold models of these loops to our cryo-EM structure (Fig. S5B-D). Intriguingly, this loop is in direct proximity to the proposed binding site for EFR3, which recruits PI4KA to the plasma membrane ^12^. In yeast, phosphorylation of Efr3 downregulates PI4KA recruitment through decreased association with the TTC7 isoform Ypp1 ^16^. Further experimentation defining the binding interface of EFR3 to the PI4KA complex should consider the possible role of these PTMs in TTC7 in regulating PI4KA membrane localisation and activity.

A important aspect in the regulation of almost all phosphoinositide kinases is their recruitment to membrane surfaces, with this mediated by multiple mechanisms, including protein-protein and protein-lipid interactions, as well as post-translational modifications ^38–40^. PI4KA is primarily active at the PM ^15^, with the CNAβ1 isoform of Calcineurin also being localised at the PM through palmitoylation ^23^. A limitation of our study is that we are studying the association of the PI4KA complex with Calcineurin in solution, and not in the biologically relevant context of the plasma membrane. There are technical challenges associated with studying a large peripheral membrane protein complex by cryo-EM and HDX-MS approaches. However, the insights gained from this work are relevant, as the presence of a membrane will likely only enhance complex formation due to increased local concentration but not alter the mode of interaction observed in our studies. Nevertheless, future studies should focus on delineating the role of PI4KA-Calcineurin interface in cellular membranes.

Overall, our work reveals important new insight into the molecular basis for how CNAβ1 can bind to and regulate the PI4KA complex. The structural insight generated by our work provides an exciting platform for continued investigations into how phosphorylation regulates PI4KA, and how calcium signaling is integrated into PI4P production at the plasma membrane.

## STAR Methods

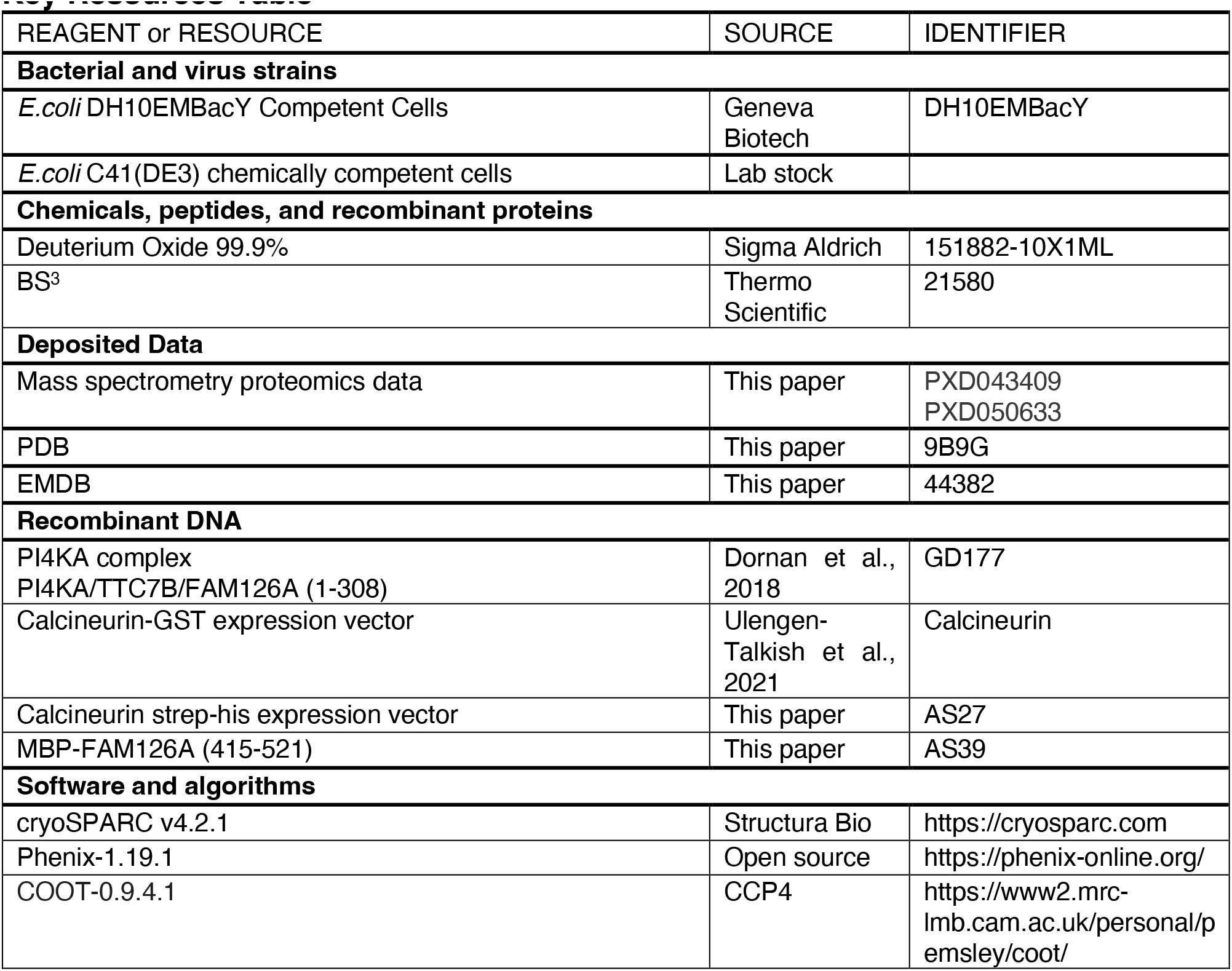
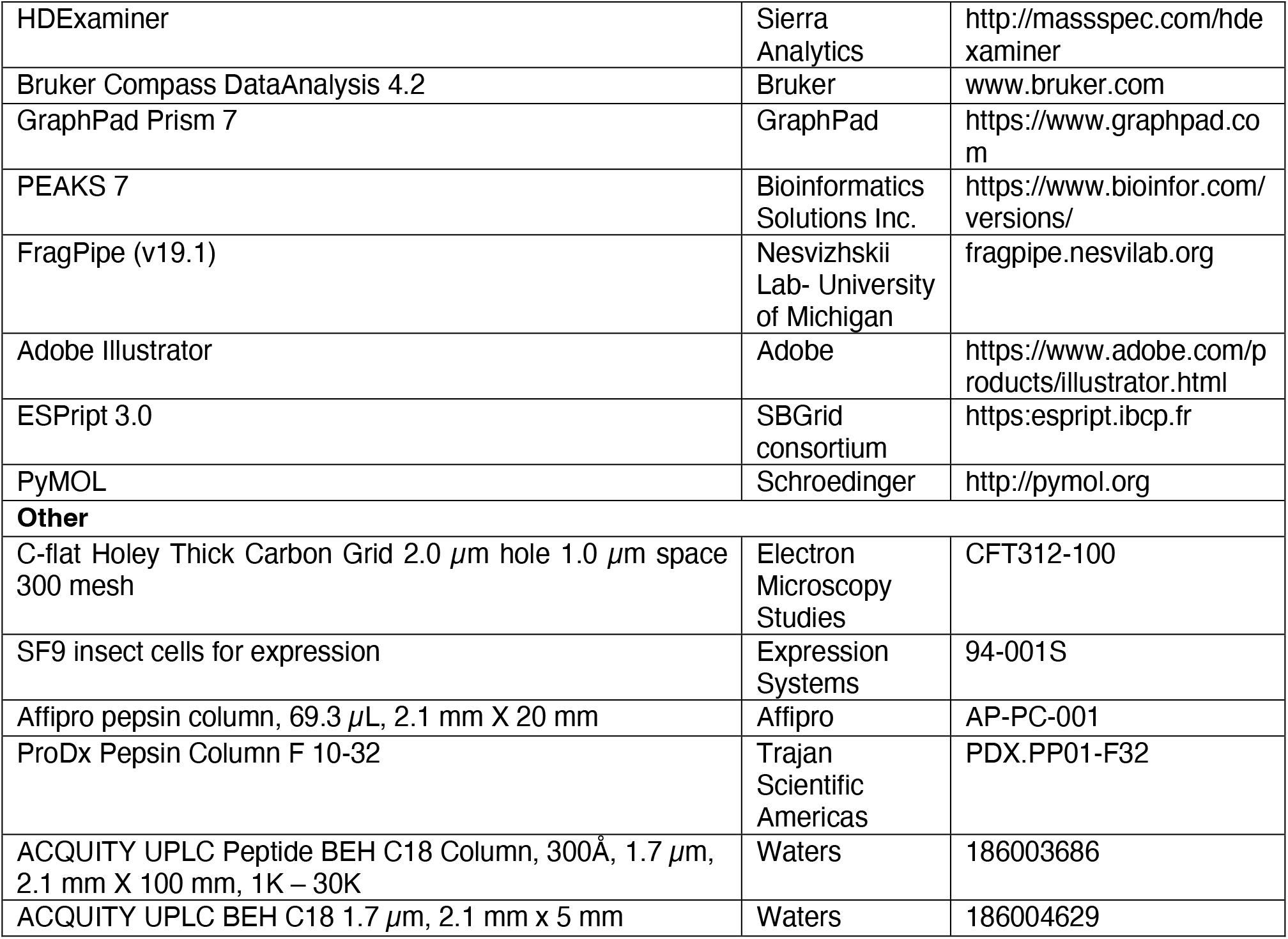
Key Resources Table.

Resource availability

## Lead contact

Further information and requests for resources and reagents should be directed to and will be fulfilled by the lead contact John E Burke

## Materials availability

All materials generated from this study will be available by request to the lead contact.

### Experimental model and subject details

PI4KA complex was expressed in *Spodoptera frugiperda* (Sf9) cells. Sf9 cells (94-001S) were obtained from Expression Systems (CA, USA) and were cultured ESF 921 media (96-001-01, Expression Systems, CA, USA) at 27°C. Calcineurin and the MBP-FAM126A construct were expressed in C41 (DE3) *E.coli* in 2xYT media.

## Method Details

### Multiple Sequence Alignments

Multiple sequence alignments were generated using Clustal Omega. Aligned sequences were then analysed by ESPript 3.0 (https://espript.ibcp.fr) ^41^ to visualize conservation. The uniprot accession codes for Figure 2C are P42356, E9Q3L2, A0A8M3AWF4, Q9W4X4, Q9XW63, Q9USR3, and P37297. The uniprot accession codes for Figure 2D are Q86TV6, Q9ULT0, E9Q6P5, Q8BGB2, A1L101, A0A8M9QA29, A0A0B4K7H0, Q7KN74, H2KYB6, O94441, and P46951. The uniprot accession codes for Figure 3G are P42356, E9Q3L2, Q49GP4, S4RUP9, ASA812BZT7, V4C7L9, Q9W4X4, Q9XW63, Q9USR3, and P37297. The uniprot accession codes for Figure 4G are Q9BYI3, Q6P9N1, Q6P121, Q8IXS8, Q8C729, A1A5W7, Q7K1C5, and Q6A586.

### Protein Expression

PI4KA complex (PI4KA/TTC7B full length, FAM126A (1-308)) was expressed from bacmids harbouring MultiBac constructs which were transfected into *Spodoptera frugiperda (Sf9) cells,* and viral stocks amplified for one generation to acquire a P2 generation viral stock. Final viral stocks were added to *Sf9 cells* in a 1/100 virus volume-to-cell ratio. Constructs were expressed for 65-72 hours before pelleting of infected cells. GST-tagged or SII-10xHis (2x strep with 10x his) human calcineurin A (aa 2-391 human CNA alpha isoform) in tandem with calcineurin B were expressed in BL21 C41 *Escherichia coli*, induced with 0.1 mM IPTG (isopropyl β-d-1-thiogalactopyranoside) and grown at 23°C overnight. SII-10xHis MBP tagged FAM126A (415-521) was expressed in BL21 C41 *Escherichia coli* induced with 0.5 mM IPTG and grown at 37°C for 3 hours. Cell pellets were washed with PBS immediately prior to being snap-frozen in liquid nitrogen, then stored at -80℃.

### Protein purification

For PI4KA complex purification, *Sf9* pellets were re-suspended in lysis buffer [100 mM NaCl, 20 mM imidazole (pH 8.0), 5% glycerol (v/v), 2 mM βMe, protease inhibitor (Protease Inhibitor Cocktail Set III, EDTA-Free)] and lysed by sonication for 2.5 minutes. Triton X-100 was added to a 0.1% (v/v) and centrifuged for 45 minutes at 20,000 g at 1℃ (Beckman Coulter J2-21, JA 20 rotor). The supernatant was loaded onto a 5 ml HisTrap FF Crude column (Cytiva), washed with NiNTA A buffer [100mM NaCl, 20 mM imidazole (pH 8.0), 5% glycerol (v/v), 2 mM βMe], washed with 10% NiNTA B buffer, and eluted with NiNTA B buffer [100 mM NaCl, 450 mM imidazole (pH 8.0), 5% glycerol (v/v), 2 mM βMe]. The eluent was loaded onto a 5 mL StrepTrap column (Cytiva) pre-equilibrated with GFB [150 mM NaCl, 20 mM imidazole (pH 7.0), 5% glycerol (v/v), 0.5 mM TCEP] then washed prior to the addition of a lipoyl domain containing TEV protease. TEV cleavage proceeded overnight before elution with 10 mL of GFB. Protein was concentrated using an Amicon 50 kDa MWCO concentrator (MilliporeSigma). PI4KA complex used for crosslinking experiments with Calcineurin was taken after this concentration step (see below, *Crosslinking*). PI4KA complex used for HDX-MS experiments was loaded onto a pre-equilibrated [150 mM NaCl, 20 mM imidazole (pH 7.0), 5% glycerol (v/v), 0.5 mM TCEP] Superose 6 Increase column (Cytiva) and protein-containing fractions were collected, concentrated, flash frozen in liquid nitrogen, and stored at -80 °C. The stoichiometry of the PI4KA based on size exclusion chromatography, SDS-PAGE, and negative stain microscopy is equimolar (2:2:2).

*Escherichia coli* cell pellets containing GST-tagged human Calcineurin were lysed by sonication for 5 minutes in lysis buffer [50 mM Tris (pH 8.0), 100 mM NaCl, 2 mM EDTA, 2 mM EGTA, protease inhibitors (Millipore Protease Inhibitor Cocktail Set III, Animal-Free)]. NaCl was then added to 1 M and the lysed solution was centrifuged at 20,000 g at 1°C for 45 minutes (Beckman Coulter J2-21, JA 20 rotor). Tween-20 was added to the supernatant to 0.1% (v/v) then loaded onto a 5 mL GSTrap 4B column (Cytiva) in a super loop fashion for 1 hour, then washed in Wash Buffer [50 mM Tris (pH 8.0), 110 mM KOAc, 2 mM MgOAc, 0.02% CHAPS (w/v), 5% glycerol (v/v) 0.5 mM TCEP] to remove non-specifically bound protein. To remove GroEL chaperone the column was washed with Wash buffer containing 2 mM ATP. To remove the GST tag, PreScission protease was added to the column and incubated overnight at 4°C. The cleaved protein was eluted with 7 mL of Wash Buffer. The eluant underwent size exclusion chromatography (SEC) using a Superdex 200 10/300 column (Cytiva) equilibrated in GFB [150 mM NaCl, 20 mM imidazole (pH 7.0), 5% glycerol (v/v), 0.5 mM TCEP] for crosslinking experiments, or Wash Buffer [50 mM Tris (pH 8.0), 110 mM KOAc, 2 mM MgOAc, 0.02% CHAPS (w/v), 5% glycerol (v/v) 0.5 mM TCEP]. For crosslinking reaction (see below, *Crosslinking*), fresh, concentrated protein was used. In all other cases, fraction(s) containing protein of interest were pooled, concentrated, flash frozen in liquid nitrogen and stored at -80°C.

*Escherichia coli* cell pellets containing SII-10xHis tagged Calcineurin were lysed by sonication for 5 minutes in lysis buffer (50 mM Tris pH 8.0 (RT), 100 mM NaCl, 2 mM EDTA, 2 mM EGTA, 10 mM Imidazole pH 8.0 (RT), 2 mM βMe, protease inhibitors (Millipore Protease Inhibitor Cocktail Set III, Animal-Free)). NaCl was added to 1 M and the lysed solution was centrifuged at 20,000 g for 50 minutes at 4°C. Tween-20 was added to the supernatant to 0.1% (v/v) then loaded onto a HisTrap FF Crude column (Cytiva), washed with NiNTA A buffer [50 mM Tris pH 8.0 (RT), 100 mM NaCl, 2 mM EDTA, 2 mM EGTA, 10 mM Imidazole pH 8.0 (RT),2 mM βMe], washed with 6% NiNTA B buffer, and eluted with NiNTA B buffer [50 mM Tris pH 8.0 (RT), 100 mM NaCl, 2 mM EDTA, 2 mM EGTA, 200 mM Imidazole pH 8.0 (RT),2 mM βMe]. The eluent was loaded onto a 5 mL StrepTrap column pre-equilibrated with GFB [20 mM HEPES (pH 7.5) RT, 150 mM NaCl, 0.5 mM TCEP] then washed prior to the addition of a lipoyl domain containing TEV protease. TEV cleavage proceeded overnight before elution with 12 mL of GFB. The eluant underwent size exclusion chromatography (SEC) using a Superdex 200 10/300 column equilibrated in GFB [150 mM NaCl, 20 mM imidazole (pH 7.0), 5% glycerol (v/v), 0.5 mM TCEP]. Fraction(s) containing protein of interest were pooled, concentrated, flash frozen in liquid nitrogen and stored at -80°C

*Escherichia coli* cell pellets containing FAM126A (415-521) were lysed by sonication for 5 minutes in lysis buffer (20 mM Tris pH 8.0 (RT), 100 mM NaCl, 20 mM Imidazole pH 8.0 (RT), 5% Glycerol (v/v), 2 mM βMe, protease inhibitors (Millipore Protease Inhibitor Cocktail Set III, Animal-Free)). The lysed solution underwent sonication for 5 minutes prior to Triton-X being added to 0.1% (v/v). The solution was then centrifuged at 20,000 g for 45 minutes at 2°C. The supernatant was then loaded onto a HisTrap FF Crude column (Cytiva), washed with High Salt buffer (20 mM Tris pH 8.0 (RT), 1 M NaCl, 20 mM Imidazole pH 8.0 (RT), 5% Glycerol (v/v), 2 mM βMe), NiNTA A buffer (20 mM Tris pH 8.0 (RT), 100 mM NaCl, 20 mM Imidazole pH 8.0 (RT), 5% Glycerol (v/v), 2 mM βMe), 6% NiNTA B, and eluted with NiNTA B (20 mM Tris pH 8.0 (RT), 100 mM NaCl, 250 mM Imidazole pH 8.0 (RT), 5% Glycerol (v/v), 2 mM βMe). The eluent was loaded onto a 5 mL StrepTrap column pre-equilibrated in GFB (20 mM HEPES pH 7.5 (RT), 150 mM NaCl, 5% Glycerol, 0.5 mM TCEP) and washed with GFB containing 0.5 M NaCl, an ATP wash (GFB with 2 mM ATP, 10 mM MgCl2, and 150 mM KCl, a GFB wash, and then eluted with GFB containing 2.5 mM desthiobiotin. Protein was concentrated and injected onto a pre-equilibrated Superdex 200 10/300 column equilibrated in GFB. Fractions containing protein were collected, aliquoted, flash frozen in liquid nitrogen, and stored at -80°C.

### PI4KA complex-Calcineurin size exclusion chromatography co-elution

PI4KA complex (1.89 µM) and Calcineurin (8.17 µM) were incubated on ice for 15 minutes prior to loading onto a Superose 6 Increase column pre-equilibrated in GFB [150 mM NaCl, 20 mM imidazole (pH 7.0), 5% glycerol (v/v), 0.5 mM TCEP]. Fractions containing the PI4KA complex bound to Calcineurin were pooled, concentrated, flash frozen in liquid nitrogen and stored at -80°C. Fresh protein was run on an SDS-PAGE gel.

### Electron Microscopy/Model Building

*Protein sample preparation using BS*^3^ *crosslinker:* PI4KA complex and Calcineurin were incubated together on ice for 15 minutes at 2 µM and 4 µM, respectively. BS3 was added to a final concentration of 1 mM and the reaction was incubated on ice for 2 hours. 1 M Tris (pH 7.5) was added to 50 mM to quench the reaction and incubated at RT for 15 minutes. The quenched reaction was loaded onto Superose 6 Increase 10/300 column in GFB [150 mM NaCl, 20 mM imidazole (pH 7.0), 5% glycerol (v/v), 0.5 mM TCEP]. Fractions encompassing the main peak, consistent with an elution volume of the complex, were collected, concentrated, flash frozen in liquid nitrogen, and stored at -80°C.

*Cryo-EM sample preparation and data collection:* C-Flat 2/1-3Cu-T-50 grids were glow-discharged for 25 s at 15 mA using a Pelco easiGlow glow discharger. 3 µl of PI4KA complex crosslinked to Calcineurin were applied to the grids at 0.77 mg/ml and vitrified using a Vitrobot Mark IV (FEI) by blotting for 1 second at a blot force of -5 at 4°C and 100% humidity. Specimens were screened using a 200-kV Glacios transmission microscope (Thermo Fisher Scientific) equipped with a Falcon 3EC direct electron detector (DED). Datasets were collected using a 300-kV Titan Krios equipped with a Falcon 4i camera and the Selectris energy filter. A total of 10,121 super-resolution movies were collected using SerialEM with a total dose of 50 e^-^/Å^2^ over 603 frames at a physical pixel size of 0.77 Å per pixel, using a defocus range of -1 to -2 um, at 165,000 x magnification.

*Cryo-EM image processing:* All data processing was carried out using cryoSPARC v4.2.1. Patch motion correction using default settings was applied to all movies to align the frames and Fourier-crop the outputs by a factor of 2. The contrast transfer function (CTF) of the resulting micrographs was estimated using the patch CTF estimation job with default settings. Micrographs were manually curated to contain only micrographs with CTF fit resolution less than or equal to 10.

To generate an initial model, 222,684 particles were picked from 2784 micrographs using blob picking with a minimum and maximum diameter of 250 and 280, respectively. Particles were inspected using the inspect picks job to remove particles that picked ice contamination and were then extracted with a box size of 480 pixels and a Fourier cropping of the output by a factor of 4, for a total of 154,266 particles. The particles were subjected to two rounds of 2D classification. A total of 43,673 particles were used for *ab*

*initio* reconstruction and heterogeneous refinement using two classes, with C1 symmetry. The best heterogeneous refinement then underwent one round of homogeneous refinement with C1 symmetry. This was used to create 2D templates using cryoSPARC’s create templates job, which were used to pick particles from the complete dataset.

To generate a cryo-EM map from the complete dataset, the above 2D templates were low pass filtered to 20 Å and used to template pick 9,830 micrographs with the particle diameter set to 450 Å. 1,181,312 particles were inspected and 604,191 were extracted with a box size of 1200 pixels and the outputs were Fourier-cropped by a factor of 2.08, and subjected to 2D classification with 40 online-EM iterations to remove particles with ice contamination or those who showed no features. 428,725 particles were used for *ab initio* reconstruction and heterogeneous refinement using two classes and C1 symmetry. 239,931 particles from the best reconstruction underwent local motion correction and a subsequent *ab initio* reconstruction and heterogeneous refinement with two classes. A total of 235,760 particles were used to carry out a homogeneous refinement with C1 symmetry using the previous 3D reconstruction as the starting model, yielding a reconstruction with an overall resolution of 3.51 Å based on the Fourier shell correlation (FSC) 0.143 criterion. Further refinement of these particles utilizing homogenous refinement, Global CTF Correction and two rounds of non-uniform refinement with C2 symmetry applied yielded a reconstruction with an overall resolution of 3.50 Å based on the Fourier shell correlation (FSC) 0.143. The full workflow to generate the final cryo-EM map is shown in Extended Figure 1.

*AlphaFold:* We used the protein prediction software, AlphaFold3 ^30^ to predict where Calcineurin interacts with both PI4KA and FAM126A.

We first modelled the PI4KA interface using the AlphaFold server (BETA) (https://golgi.sandbox.google.com/) and input the sequence of our PI4KA/TTC7B/FAM126A ΔC complex construct and our Calcineurin construct (CNA alpha 1 (2-391) and CNB) with cofactors (Fe^2+^, 4 x Ca^2+^). The resulting top ranked model had ptm and iptm scores of 0.76 and 0.74, respectively, consistent with a stable complex. To evaluate the confidence of individual subunit assembly predictions, based on our previous knowledge from previously published X-ray and Cryo-EM structure of TTC7B/FAM126A and PI4KA/TTC7B/FAM126A, respectively, we analyzed the chain_pair_iptm and chain_pair_pae_min values. The chain_pair_iptm scores are useful in evaluating the confidence of predicted protein-protein interfaces. The chain_pair_iptm score for PI4KA:TTC7B was 0.84, TTC7B:FAM126A was 0.89, PI4KA:CNA was 0.72, PI4KA:CNB was 0.54 and CNA:Fe^2+^ was 0.91. The chain_pair_pae_min values correlate with whether two chain interact with each other. The chain_pair_pae_min score for PI4KA:CNA was 3.39, and PI4KA:CNB was 11.24. The predicated aligned error (PAE) for the top-ranked output is shown in Figure 3B. These metrics correlate with previously published structural information and our Cryo-EM and HDX-MS data therefore we were confident with this predicted model.

To predict the interaction between Calcineurin and FAM126A’s C-terminus, we used Alphafold3 with a search composed of the C-terminus of FAM126A (aa 390-521) with Calcineurin (CNA alpha: aa 2-391, and CNB) with cofactors (4 x Ca^2+^). The resulting top ranked model had ptm and iptm scores of 0.81 and 0.76. The predicted aligned error (PAE) for the top-ranked output is shown in Figure 5B. The chain_pair_iptm score for FAM126ACterm:CNA was 0.74 and for FAM126ACterm:CNB was 0.63. The chain_pair_pae_min score for FAM126ACterm:CNA was 2.39, and FAM126ACterm:CNB was 3.85.

*Model building:* The cryo-EM structure of PI4KA-TTC7B-FAM126A (PDB: 6BQ1) ^10^ was fit into the map using Chimera ^42^. The horn domain of PI4KA was at higher local resolution than the original data, and allowed for iterative rounds of automated model building in Phenix guided by secondary structure predicted by AlphaFold ^24^, manual model building in COOT, and refinement in Phenix.real_space_refine using realspace, rigid body, and adp refinement with tight secondary structure restraints ^43^. This allowed for building the full solenoid horn domain with high confidence.

The local resolution of Calcineurin was the lowest of any protein chains, although the high resolution structure of Calcineurin (PDB: 6NUC) ^26^ could be fit into the density. We then used the AlphaFold3 search result for Calcineurin bound to the beta strand of PI4KA (see previous section), and fit into the map using Chimera ^42^. The entire model was refined using Phenix.real_space_refine using rigid body, adp refinement with both reference model restraints using the high resolution Calcineurin model 6NUC and tight secondary structure restraints ^43^. The full refinement and validation statistics are shown in Supplementary Table 1.

### HDX-MS

*HDX-MS sample preparation:* Initial HDX reactions comparing PI4KA complex apo and Calcineurin apo to PI4KA complex + Calcineurin were carried out in an 8.16 µl reaction volume containing 11 pmol of both proteins. The exchange reactions were initiated by the addition of 5.88 µL of D2O buffer (20 mM Imidazole pH 7, 150 mM NaCl, 93.9% D2O (V/V)) to 2.88 µL of protein (final D2O concentration of 67.7%). Reactions proceeded for 3s, 30s, and 300s at room temperature (18°C) before being quenched with ice cold acidic quench buffer, resulting in a final concentration of 0.6M guanidine HCl and 0.9% formic acid post quench. HDX reactions comparing PI4KA complex apo to PI4KA complex + Calcineurin were then carried out at pH 6.5 to capture changes in disordered and flexible regions of the PI4KA complex. Reactions occurred in an 8.12 µl reaction volume containing 10 pmol of PI4KA complex and 20 pmol of Calcineurin. The exchange reactions were initiated by the addition of 5.74 µL of D2O buffer (20 mM MES pH 6.5, 150 mM NaCl, 1 mM CaCl2, 93.37% D2O (V/V)) to 2.38 µL of protein (final D2O concentration of 67%). Reactions proceeded for 3s, 10s, and 30s at 0°C before being quenched with ice cold acidic quench buffer, resulting in a final concentration of 0.6M guanidine HCl and 0.9% formic acid post quench. All conditions and time points were created and run in independent triplicate. Samples were flash frozen immediately after quenching and stored at -80°C.

HDX reactions comparing FAM126A (415-521) apo to FAM126A (415-521) + Calcineurin were carried out in a 12 µl reaction volume containing 30 pmol of both proteins. The exchange reactions were initiated by the addition of 9 µL of D2O buffer (20 mM MES pH 6.5, 150 mM NaCl, 1mM CaCl2, 93.4% D2O (V/V)) to 3 µL of protein (final D2O concentration of 70%). Reactions proceeded for 3s, 10s, and 30s at 0°C, pH 6.5 before being quenched with ice cold acidic quench buffer, resulting in a final concentration of 0.6M guanidine HCl and 0.9% formic acid post quench. All conditions and time points were created and run in independent triplicate. A fully deuterated sample was made according to the protocol of ^44^ by incubating 1 µl of 30 µM FAM126A (415-521) with 7 M guanidine HCl (final 4.67M), heated to 90°C for five minutes followed by two minutes 20°C, followed by the addition of 9 µl of D2O buffer (20 mM MES pH 6.5, 150 mM NaCl, 1 mM CaCl2, 93.37% D2O (V/V)) for 10 minutes at 50°C then two minutes at 20°C, and then 0°C for 2 minutes before being quenched as above. Samples were flash frozen immediately after quenching and stored at -80°C.

*Protein digestion and MS/MS data collection:* Protein samples were rapidly thawed and injected onto an integrated fluidics system containing a HDx-3 PAL liquid handling robot and climate-controlled (2°C) chromatography system (Trajan), a Dionex Ultimate 3000 UHPLC system (Thermofisher), or a Acquity UPLC I-Class Series System (Waters), as well as an Impact HD QTOF Mass spectrometer (Bruker). The full details of the automated LC system are described in ^28^. Samples comparing PI4KA complex apo and Calcineurin apo to PI4KA complex + Calcineurin (pH 7.0, 18°C) were run over one immobilized pepsin column (Trajan; ProDx protease column, 2.1 mm x 30 mm, PDX.PP01-F32) at 200 µL/min for 3 minutes at 10°C. Samples comparing FAM126A (415-521) apo to FAM126A (415- 521) + Calcineurin and samples comparing PI4KA complex apo to PI4KA complex + Calcineurin (pH 6.5, 0°C) were run over an immobilized pepsin column (Affipro; AP-PC- 001) at 200 µL/min for 4 minutes at 2°C. The resulting peptides were collected and desalted on a C18 trap column (ACQUITY UPLC BEH C18 1.7 µm column, 2.1 mm x 5 mm; Waters 186004629). The trap was subsequently eluted in line with an ACQUITY 1.7 µm particle, 2.1 mm × 100 mm C18 UPLC column (Waters; 186003686), using a gradient of 3-35% B (Buffer A 0.1% formic acid; Buffer B 100% acetonitrile) over 11 minutes immediately followed by a gradient of 35-80% over 5 minutes. Mass spectrometry experiments were acquired over a mass range from 150 to 2200 m/z using an electrospray ionization source operated at a temperature of 200°C and a spray voltage of 4.5 kV.

*Peptide identification:* Peptides were identified from the non-deuterated samples of the PI4KA complex, Calcineurin, or FAM126A (415-521) using data-dependent acquisition following tandem MS/MS experiments (0.5 s precursor scan from 150-2000 m/z; twelve 0.25 s fragment scans from 150-2000 m/z). For HDX reactions comparing PI4KA complex apo and Calcineurin apo to PI4KA complex + Calcineurin (pH 7.0, 18°C), MS/MS datasets were analysed using PEAKS7 (PEAKS), and peptide identification was carried out by using a false discovery based approach, with a threshold set to 0.1% using a database of purified proteins and known contaminants ^45^. The search parameters were set with a precursor tolerance of 20 ppm, fragment mass error 0.02 Da, charge states from 1-8, with a -10logP score of 31.9 and 15 for the PI4KA complex or Calcineurin respectively.

For HDX reactions comparing PI4KA complex apo to PI4KA complex + Calcineurin at pH 6.5, and FAM126A (415-521) apo to FAM126A (415-521) in complex with Calcineurin, MS/MS datasets were analysed using FragPipe v18.0 and peptide identification was carried out by using a false discovery-based approach using a database of purified proteins and known contaminants ^46–48^. MSFragger was utilised, and the precursor mass tolerance error was set to -20 to 20 ppm. The fragment mass tolerance was set at 20 ppm. Protein digestion was set as nonspecific, searching between lengths of 4 and 50 aa, with a mass range of 400 to 5000 Da. For the Sf9 expressed PI4KA complex the search was carried out with variable phosphorylation of S, T, and Y. Phosphorylated sites in the PI4KA complex were identified in PI4KA between residues 252-281 and in TTC7B residues 153-164, with these sites being 35% and 50% percent phosphorylated, respectively.

## Quantification and statistical analysis

### Mass Analysis of Peptide Centroids and Measurement of Deuterium Incorporation

HD-Examiner Software (Sierra Analytics) was used to automatically calculate the level of deuterium incorporation into each peptide. All peptides were manually inspected for correct charge state, correct retention time, appropriate selection of isotopic distribution, etc. Deuteration levels were calculated using the centroid of the experimental isotope clusters. Results are presented as relative levels of deuterium incorporation and the only control for back exchange in the PI4KA complex containing experiments was the level of deuterium present in the buffer (67.7% and 67%). For FAM126A (415-521) HDX experiments, back exchange was controlled for by generating a fully deuterated sample. For all HDX-MS experiments, differences in exchange in a peptide were considered significant if they met all three of the following criteria: ≥5% change in exchange, ≥0.45 Da difference in exchange, and a p value <0.01 using a two tailed student t-test. The raw HDX data are shown in two different formats. The raw peptide deuterium incorporation graphs for a selection of peptides with significant differences are shown in Figures 3C/4D/S4B-E/S4G, with the raw data for all analysed peptides in the source data. To allow for visualization of differences across all peptides, we utilised number of deuteron difference (#D) plots (Fig. 4E/5E/S4A/S4F). These plots show the total difference in deuterium incorporation over the entire H/D exchange time course, with each point indicating a single peptide. Samples were only compared within a single experiment and were never compared to experiments completed at a different time with a different final D2O level. The data analysis statistics for all HDX-MS experiments are in the source data file according to the guidelines of ^49^. The mass spectrometry proteomics data have been deposited to the ProteomeXchange Consortium via the PRIDE partner repository ^50^ with the dataset identifiers PXD043409 and PXD050633.

## Data and code availability

- The EM data have been deposited in the EM data bank with accession number (EMDB: 44382), and the associated structural model has been deposited to the PDB with accession number (PDB: 9B9G). The MS proteomics data have been deposited to the ProteomeXchange Consortium via the PRIDE ^50^ partner repository with the dataset identifiers PXD043409 and PXD050633. All raw data in all figures are available in the source data excel file.
- This paper does not report original code.
- Any additional information required to reanalyze the data reported in this paper is available from the lead contact upon request.

## Acknowledgements

JEB is supported by the Natural Science and Engineering Research council (Discovery grant NSERC-2020-04241), and a Michael Smith Foundation for Health Research (Scholar Award 17686). ALS was supported by an NSERC CGS-M scholarship. C.K.Y. is supported by CIHR (PJT-168907) and the Natural Sciences and Engineering Research Council of Canada (RGPIN-2018-03951). Cryo-EM specimens were prepared, and data was collected at the High Resolution Macromolecular Electron Microscopy (HRMEM) facility at the University of British Columbia (https://cryoem.med.ubc.ca). We thank Claire Atkinson, Joeseph Felt, Liam Worrall and Natalie Strynadka. HRMEM is funded by the Canadian Foundation for Innovation and the British Columbia Knowledge Development Fund. We thank Dr. Martha Cyert for their insight and suggestions in the preparation of the manuscript and for gifting us the GST Calcineurin expression vector.

## Author contributions

Conceptualization, A.L.S., S.S., and J.E.B.; data curation, A,L.S., S.S., MLJ and M.A.H.P.; formal analysis A.L.S., S.S., M.A.H.P., M.L.J., and J.E.B.; investigation A.L.S., S.S., M.A.H.P., N.J.H., M.L.J, and J.E.B.; visualization, A.L.S., S.S. and JEB; writing – original draft, A.L.S., S.S., and J.E.B.; writing – reviewing and editing, A.L.S., S.S., M.A.H.P., N.J.H., M.L.J., C.K.Y., and J.E.B.; methodology, A.L.S., M.A.H.P, M.L.J., C.K.Y., and J.E.B.; supervision, C.K.Y., and J.E.B.; funding acquisition, C.K.Y., and J.E.B.; project administration, J.E.B.

## Supplemental 986 Figures Tables

**Table S1.**
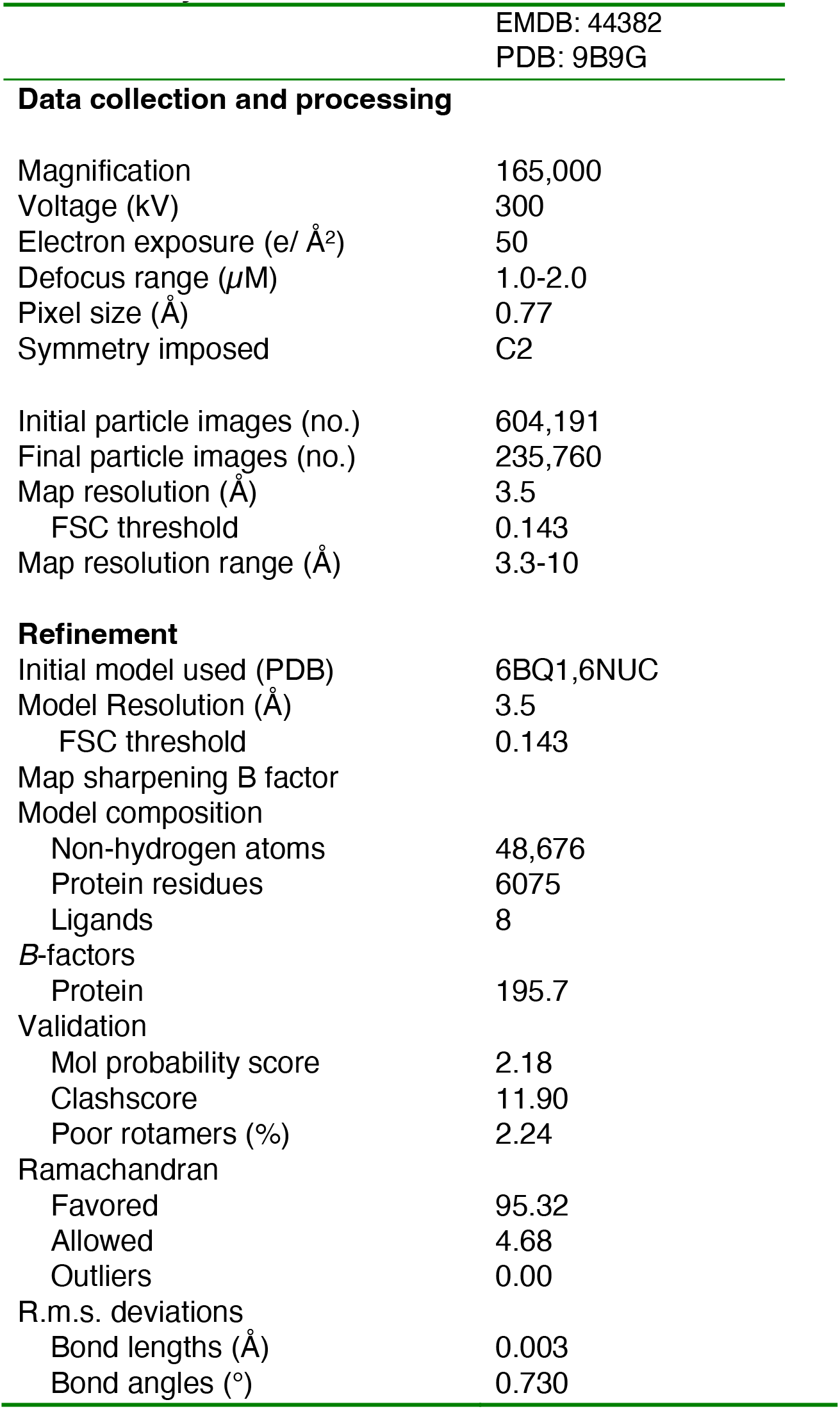
Cryo-EM data collection, refinement, and validation statistics

**Data collection and processing**
|  |  |
| --- | --- |
| Magnification | 165,000 |
| Voltage (kV) | 300 |
| Electron exposure (e/ Å<sup>2</sup>) | 50 |
| Defocus range (μM) | 1.0-2.0 |
| Pixel size (Å) | 0.77 |
| Symmetry imposed | C2 |
| Initial particle images (no.) | 604,191 |
| Final particle images (no.) | 235,760 |
| Map resolution (Å) | 3.5 |
| FSC threshold | 0.143 |
| Map resolution range (Å) | 3.3-10 |
| **Refinement** |  |
| Initial model used (PDB) | 6BQ1,6NUC |
| Model Resolution (Å) | 3.5 |
| FSC threshold | 0.143 |
| Non-hydrogen atoms | 48,676 |
| Protein residues | 6075 |
| Ligands | 8 |
| *B*-factors |  |
| Protein | 195.7 |
| Validation |  |
| Mol probability score | 2.18 |
| Clashscore | 11.90 |
| Poor rotamers (%) | 2.24 |
| Ramachandran |  |
| Favored | 95.32 |
| Allowed | 4.68 |
| Outliers | 0.00 |
| R.m.s. deviations |  |
| Bond lengths (Å) | 0.003 |
| Bond angles (°) | 0.730 |

**Supplemental Figure 1.**
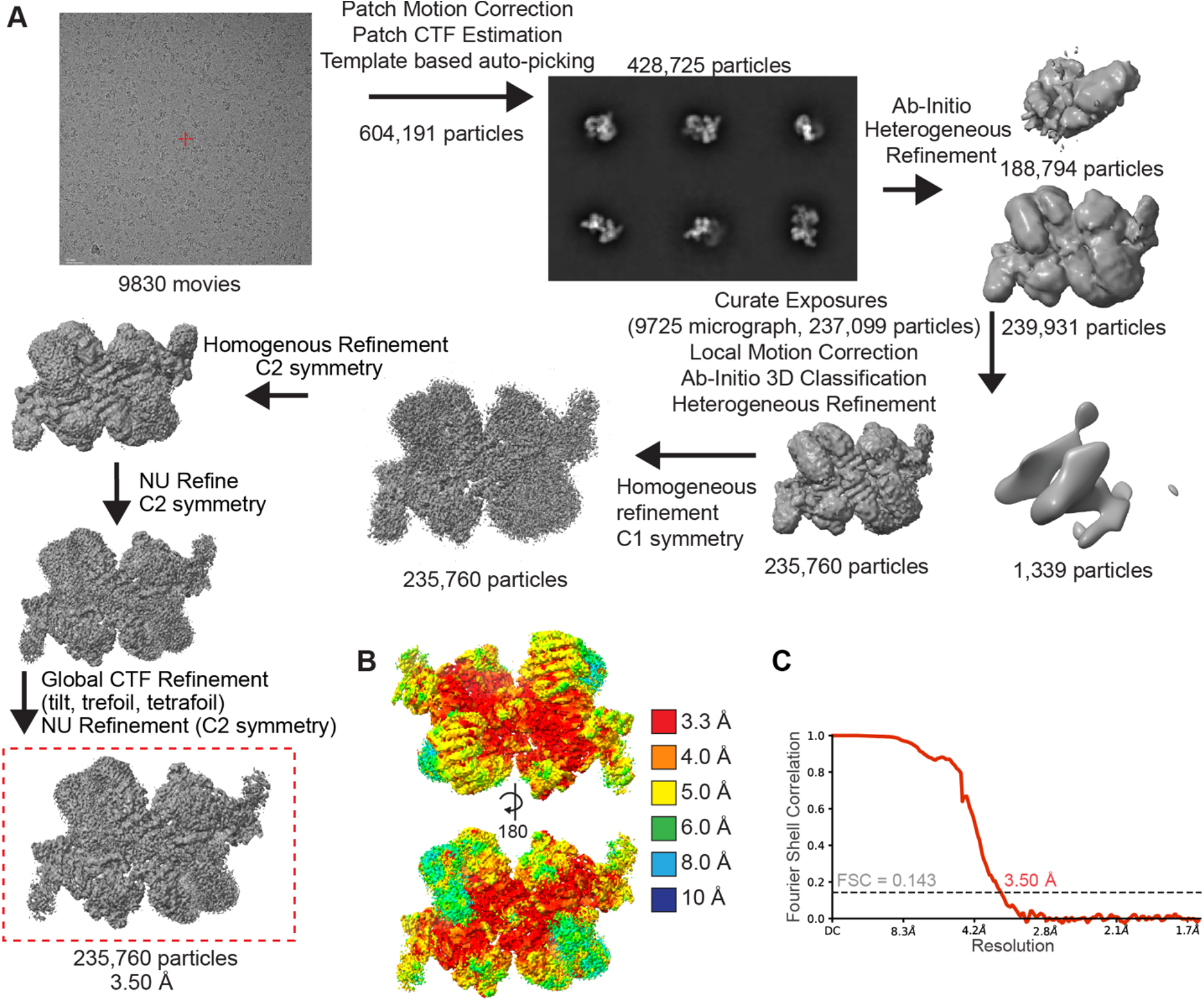
Cryo-EM data processing **A.** Cryo-EM data processing workflow showing a representative micrograph from screening on the 200 kV Glacios, representative 2D class averages, and the image processing strategy used to generate a 3D reconstruction of the PI4KA/TTC7B/FAM126A/Calcineurin complex. **B.** Final PI4KA/TTC7B/FAM126A/ Calcineurin map coloured according to local resolution estimated using cryoSPARC v4.2.1 (FSC threshold 0.143) **C.** Gold standard Fourier shell correlation coefficient (FSC) curve after auto tightening by cryoSPARC for the PI4KA/TTC7B/FAM126A/ Calcineurin map.

**Supplemental Figure 2.**
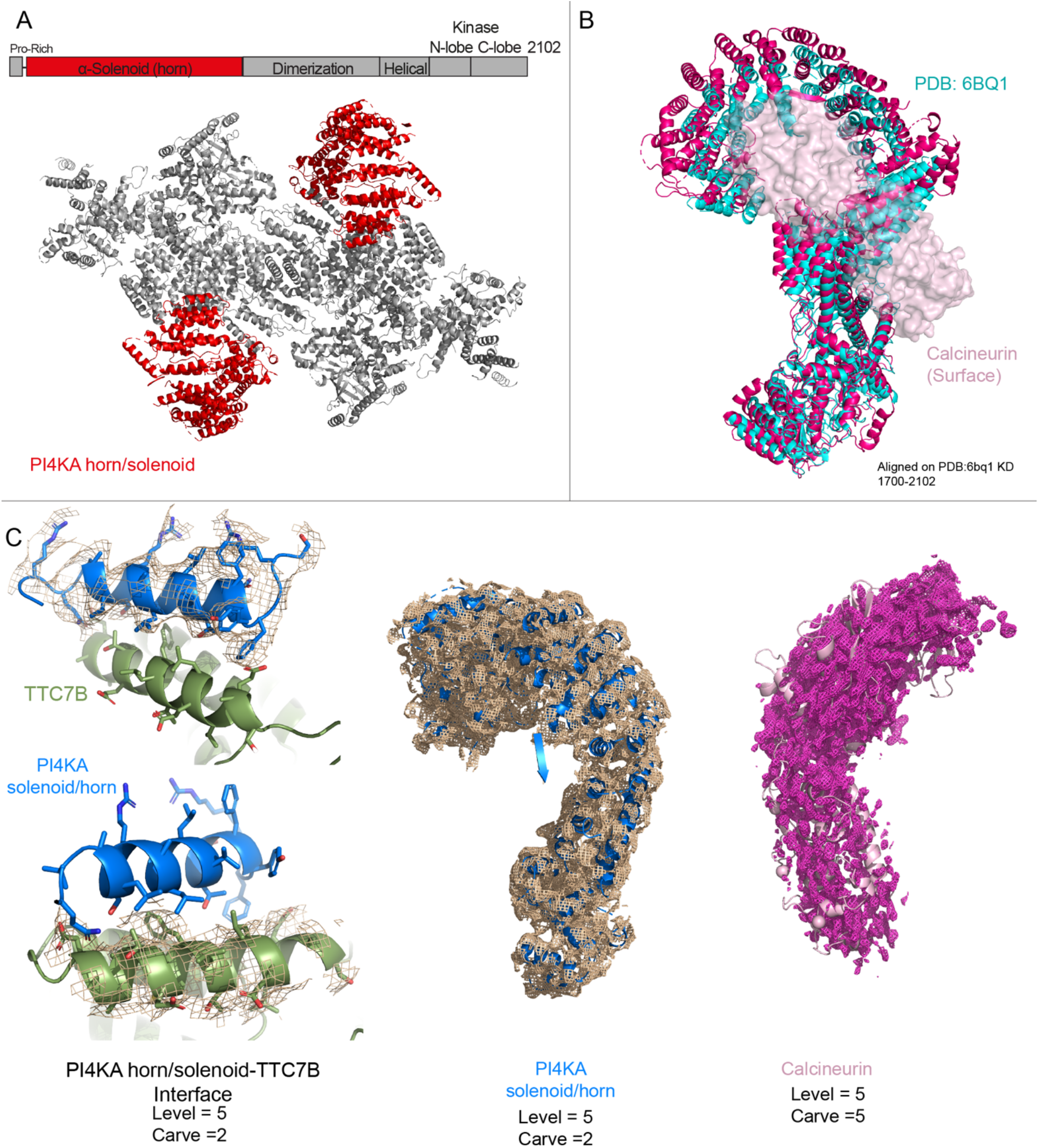
Model density fit **A.** Unique features in the PI4KA complex resolved from the cryo-EM map of the PI4KA complex interacting with Calcineurin. Red areas represent unique features, annotated on the domain schematic, and mapped onto the structural model with Calcineurin removed. **B.** Comparison of the PI4KA structure from our study (magenta) to PI4KA from PI4KA/TTC7B/FAM126A (2-289) (cyan) apo (PDB:6BQ1) (Lees *et al*, 2017). Our model was aligned to 6BQ1 using residues 1700- 2102. **C.** Electron density of selected regions. (L) TTC7B-PI4KA solenoid/horn interface, with density for the N-terminal helix of PI4KA shown on top, and the density for the TTC7B interface helix shown on bottom (M) PI4KA solenoid/horn (aa 27-929), and (R) Calcineurin.

**Supplemental Figure 3.**
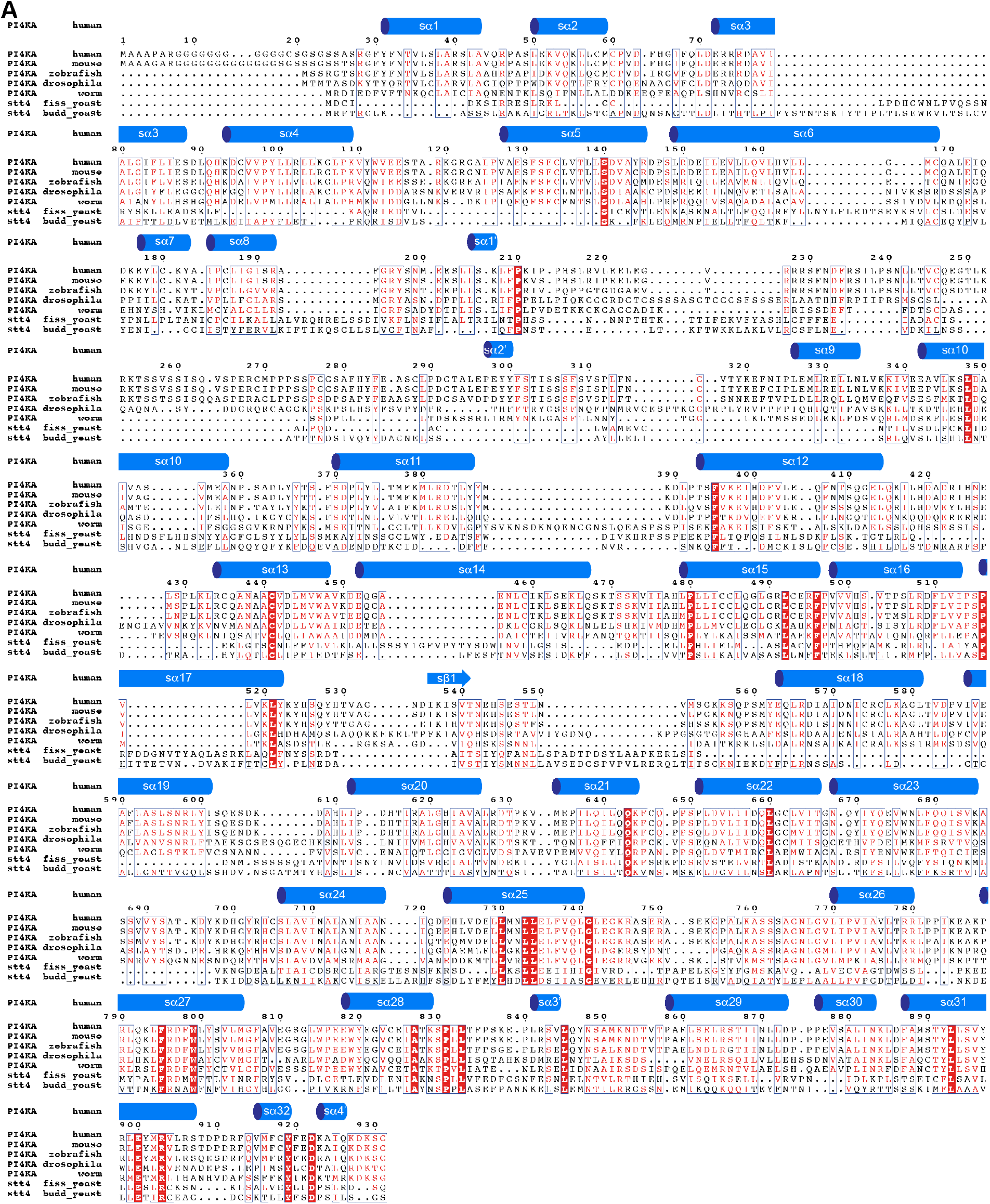
Secondary structure annotation of PI4KA solenoid/horn **A.** Multiple sequence alignment (generated with ESPript 3.0) of PI4KA from *Homo sapiens, Mus musculus, Danio rerio, Drosophila melanogaster, Caenorhabditis elegans*, *Schizosaccharomyces pombe,* and *Saccharomyces cerevisiae.* Secondary structure elements from the human PI4KA horn region, as determined from the cryo-EM map of the PI4KA complex bound to Calcineurin, are illustrated above the alignment.

**Supplemental Figure 4.**
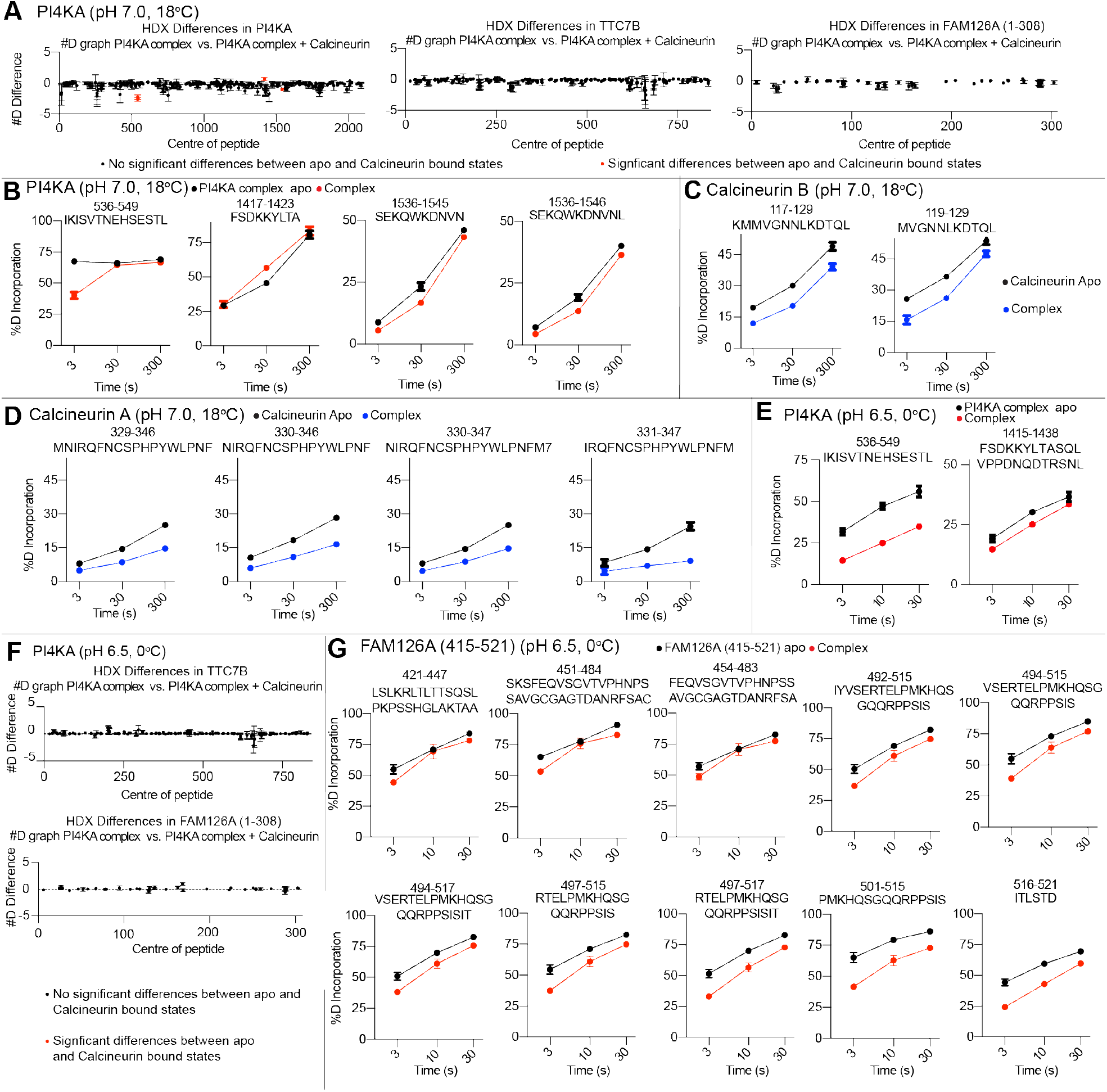
HDX-MS data underlying all experiments **A.** Sum of the number of deuteron difference of (L) PI4KA, (M) TTC7B, and (R) FAM126A (1-308) upon complex formation with Calcineurin analysed over the entire deuterium exchange time course (pH 7.0, 18°C). Each point is representative of the centre residue of an individual peptide. Peptides that met the significance criteria (see panels **B**) are colored red. Error is shown as standard deviation (SD) (n=3). The full source data underlying all HDX-MS data is included as an excel file. **B.** Selected deuterium exchange time courses of PI4KA peptides (pH 7.0, 18°C) that showed significant (defined as >5% 0.45 Da, and p<0.01 in an unpaired two-tailed t test at any time point) changes in exchange. Error is shown as standard deviation (SD) (n=3). **C.** Selected deuterium exchange time courses of Calcineurin B peptides (pH 7.0, 18°C) that showed significant (defined as >5% 0.45 Da, and p<0.01 in an unpaired two-tailed t test at any time point) decreases in exchange. Error is shown as standard deviation (SD) (n=3). **D.** Selected deuterium exchange time courses of Calcineurin A peptides (pH 7.0, 18°C) that showed significant (defined as >5% 0.45 Da, and p<0.01 in an unpaired two-tailed t test at any time point) decreases in exchange. Error is shown as standard deviation (SD) (n=3). **E.** Selected deuterium exchange time courses of PI4KA peptides (pH 6.5, 0°C) that showed significant (defined as >5% 0.45 Da, and p<0.01 in an unpaired two-tailed t test at any time point) decreases or increases in exchange. Error is shown as standard deviation (SD) (n=3). **F.** Sum of the number of deuteron difference of (L)TTC7B, and (R) FAM126A (1-308) upon complex formation with Calcineurin analysed over the entire deuterium exchange time course (pH 6.5, 0°C). Each point is representative of the centre residue of an individual peptide. Peptides that met the significance criteria (see panels **E**) are colored red. Error is shown as standard deviation (SD) (n=3). The full source data underlying all HDX-MS data is included as an excel file. **G.** Selected deuterium exchange time courses FAM 126A (415-521) peptides (pH 6.5, 0°C) that showed significant (defined as >5% 0.45 Da, and p<0.01 in an unpaired two-tailed t test at any time point) decreases or increases in exchange. Error is shown as standard deviation (SD) (n=3).

**Supplemental Figure 5.**
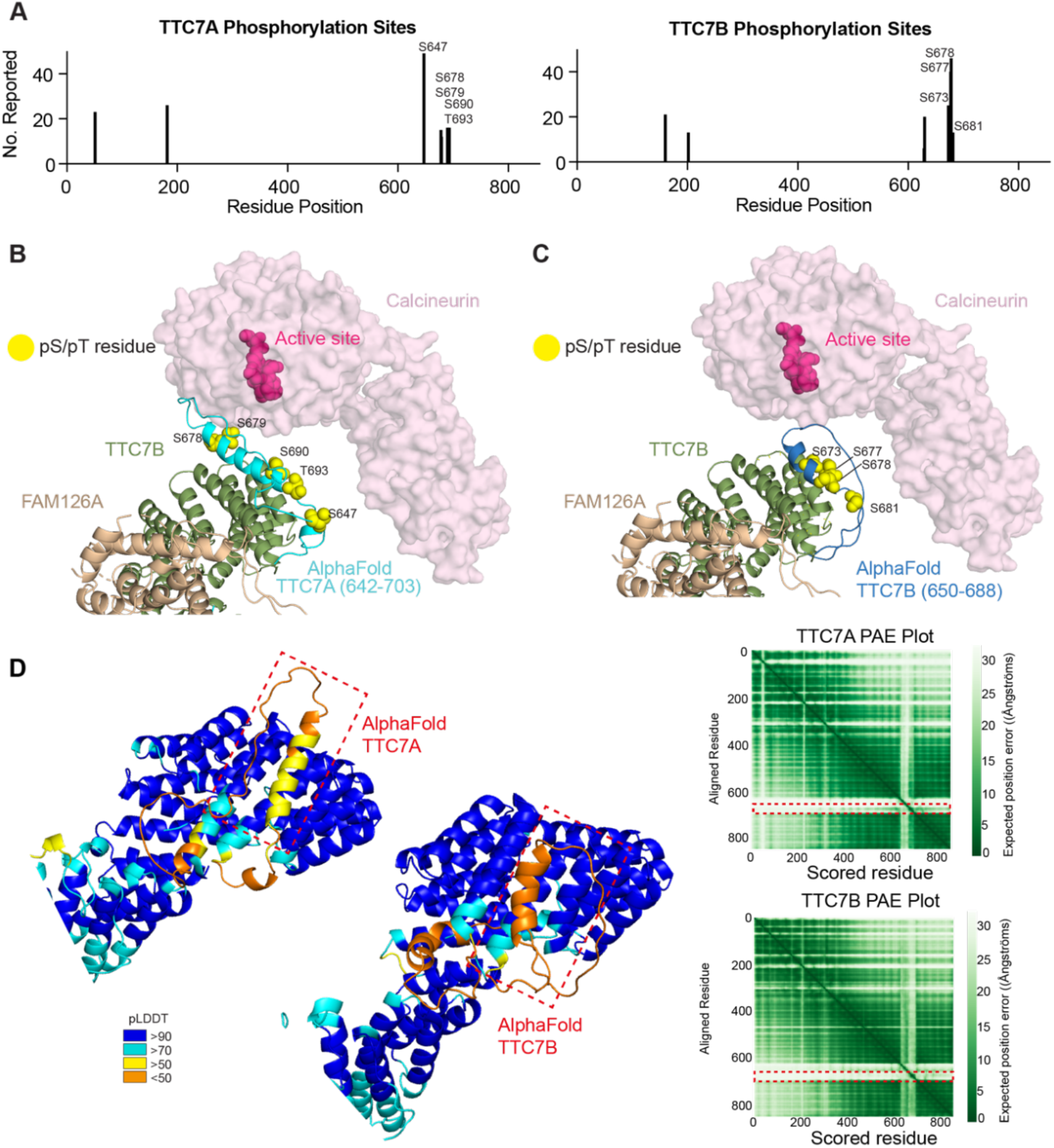
AlphaFold modelling of TTC7A/B on cryo-EM structure reveals evolutionary conserved phosphorylation sites in TTC7A and TTC7B in close proximity to Calcineurin active site. **A.** Phosphorylated serine/threonine residues in TTC7A (left) and TTC7B (right) reported on PhosphoSite (Hornbeck *et al*, 2015). Serine/threonine residues reported more than five times are shown. **B.** Multiple reported TTC7A phosphorylation sites are in proximity to the Calcineurin active site. AlphaFold model of the disordered loop in TTC7A (642-703) is aligned to the experimental density for TTC7B. Threonine/serine residues reported to be phosphorylated are shown as yellow spheres. The AlphaFold model used for the alignment are shown in panel D. **C.** Multiple reported TTC7B phosphorylation sites are in proximity to the Calcineurin active site. AlphaFold model of the disordered loop in TTC7B (650-688) is aligned to the experimental density for TTC7B. Threonine/serine residues reported to be phosphorylated are shown as yellow spheres. The AlphaFold model used for the alignment are shown in panel D. **D.** (L) AlphaFold models of TTC7A and TTC7B coloured based on pLDDT according to the legend. The disordered loop not present in the experimental structure is boxed in red. The predicted alignment error (PAE) for both AlphaFold models are shown, with the areas boxed in red indicating the disordered loop in close proximity to Calcineurin in the PI4KA-TTC7-FAM126-Calcineurin structure.

